# Comparative connectomics of parasite esophagus suggests evolutionary simplification of a nervous system

**DOI:** 10.64898/2026.07.28.741281

**Authors:** Jaeyeong Han, Abby R. Thompson, Ellie Conklin, Lav R. Varshney, Nathan E. Schroeder

## Abstract

Several prominent examples suggest that the evolution of parasitism is accompanied by nervous system simplification. However, it is unclear if this is a generalizable rule and whether parasite-associated simplification occurs at the level of synaptic connectivity. The nematode esophagus is a specialized neuromuscular feeding organ that varies with diet and lifestyle. Plant-parasitic nematodes are major agricultural pests that feed through a protrusible stylet and release of extensive glandular effectors; however, the neuronal mechanisms controlling parasite feeding are unclear. Here, we used serial-section electron microscopy to reconstruct the esophageal connectome of the infective second-stage juvenile of the soybean cyst nematode, *Heterodera glycines*, and compared it with those of the free-living species *Caenorhabditis elegans* and *Pristionchus pacificus*. Similar to these species, *H. glycines* has 20 esophageal neurons with relatively conserved cell body positions. Despite this conservation, the *H. glycines* chemical synaptic network is highly reduced in output to musculature. A unique ensheathment of neurons by a gland cell facilitates novel synaptic connectivity in *H. glycines*. The *H. glycines* esophageal network is strongly biased toward monadic synapses and shows a greater proportion of neuron-neuron and neuron-gland connections. Consistent with a reduction in motor output, network analysis indicates that the *H. glycines* esophageal network is smaller and less clustered than free-living species. Using centrality analysis and synthetic ablation, we predict that control of multiple feeding modules in *H. glycines* depends on distinct neurons compared to free-living species. These findings show how parasitism reshapes a feeding circuit and identify candidate species-specific circuits for parasite control.

**Significance statement:** Animal nervous systems are tuned to the behaviors they support. Nematodes occupy diverse ecological niches, and their feeding organ, the esophagus, is specialized to their diet. Understanding the esophageal nervous system provides evolutionary insights into parasitism as well as pathways for future control targets. We present the esophageal connectome of the soybean cyst nematode *Heterodera glycines*, which causes devastating damage to soybean production worldwide. By comparing *H. glycines* and free-living species, which diverged over 350 million years ago, we identify evolutionarily conserved features of the feeding circuit and highlight parasite-specific circuits as candidate targets for control.

## Introduction

The transition from a free-living to parasitic lifestyle is often accompanied by changes in the nervous system structure and function necessary for specialized infection behaviors. In several instances, the evolution of parasitism is correlated with nervous system simplification (1–3).

Despite advances in volumetric electron microscopy (EM), and the description of multiple connectomes from diverse organisms, a complete synaptic-level analysis of changes during the evolution of parasitism is unavailable (4–7).

Nematodes are well suited for comparative connectomics as cellular identity is highly conserved across deeply diverged species (4, 8–10). The free-living (non-parasitic) nematodes *Caenorhabditis elegans* and *Pristionchus pacificus* diverged approximately 290 million years ago (11) and have been the target of several connectomics studies (4, 9, 12).

Parasitism has evolved multiple times within the phylum Nematoda and is associated with specialized behavioral responses to host cues (13–15). The evolution of plant-parasitism among nematodes is associated with extensive morphological and functional adaptations to the esophagus. *C. elegans* uses a near simultaneous wave of esophageal muscle contraction to ingest bacteria (16). In contrast, plant-parasitic nematodes use morphologically distinct and spatially separated muscle classes, which contract independently, but in a coordinated fashion (**Fig. 1A**) (14). The soybean cyst nematode, *Heterodera glycines,* is a major agricultural pest of soybean worldwide which diverged over 350 million years ago from the common ancestor of *C. elegans* and *P. pacificus* (**Fig. S1A and B**) (11). Previous EM studies described the esophageal structure of *H. glycines* (17, 18). However, these studies did not include complete series, thereby precluding comprehensive reconstructions and systematic annotation of synaptic connectivity.

**Fig. 1.**
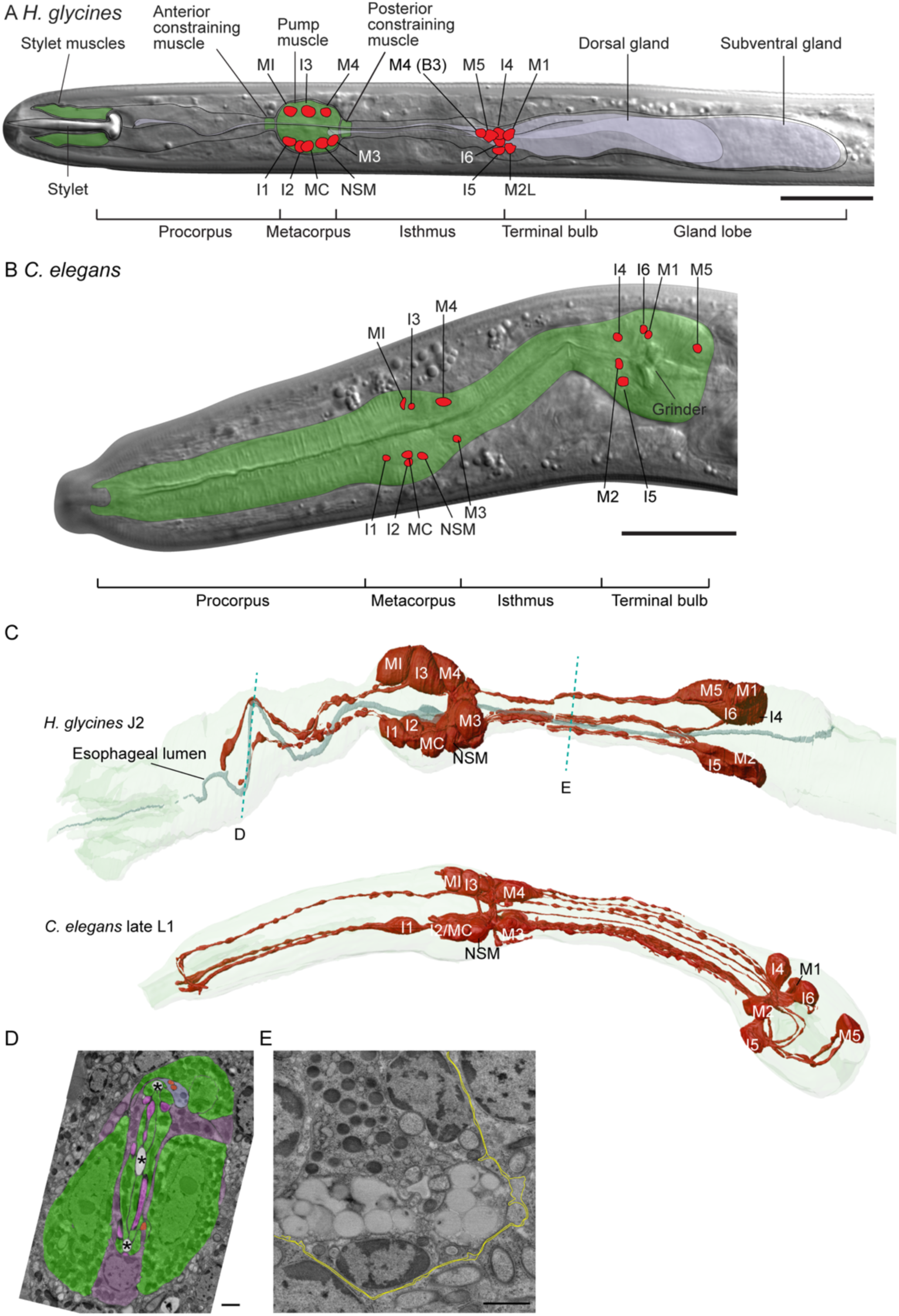
Anatomical differences between *H. glycines* and *C. elegans*. (A) DIC micrograph of J2 *H. glycines* anterior highlighting specific tissue types and positions of neuronal nuclei (red). The esophagus musculature (contractile areas in green) consists of three major types: stylet- associated muscles, anterior and posterior constraining muscles, and metacorpus pump muscles. Plant-parasitic nematode gland cells (lavender) have greatly enlarged during evolution to promote parasitism. The M4 cell body position shows variation between datasets F3 and B3. For clarity, only one nucleus for bilateral pairs is shown. (B) DIC micrograph of L2 *C. elegans* esophagus. In contrast to *H. glycines* nearly the entire esophagus of *C. elegans* is lined with radial muscle fibers and esophageal gland cells are enclosed within the terminal bulb. (C) Three-dimensional reconstruction of the esophageal nervous system in *H. glycines* J2 and *C. elegans* late L1 (lateral views). In *H. glycines,* the esophageal lumen meanders within the procorpus. Dashed lines indicate approximate locations of transverse EM sections (D, E). (D, E) Transverse EM images showing asymmetry within the *H. glycines* procorpus (D) and reduced extracellular matrix (E) (basement membrane and pseudocoelom, yellow). Scale bars: 20 µm (A) and 1 µm (D, E).

Here, we reconstruct the complete esophageal connectome of the infective second stage juvenile (J2) of *H. glycines,* using serial-section EM and compare network properties with the established adult esophageal connectomes of free-living *C. elegans* and *P. pacificus*. To account for developmental stage differences, we further reconstructed the esophageal connectome of a late L1 *C. elegans* from a previously published EM dataset (19). Our data suggests several esophageal nervous system adaptations and potential cellular targets for control of plant-parasitic nematodes.

## Results

### Conserved overall structure of the *H. glycines* esophageal nervous system with species-specific divergence

We reconstructed the esophageal connectome of two J2 *H. glycines* individuals (B3 and F3) (**Fig. S1C and D**). Like *C. elegans* and *P. pacificus,* we identified 20 esophageal neurons in *H. glycines* (**Fig. 1A**). These are spatially divided into two clusters found in the metacorpus and the posterior terminal bulb. The *H. glycines* metacorpus neurons are arranged in two subventral rows of five and one dorsal row of three, a pattern typical of other nematodes (8, 9, 17, 18, 20–22). This topological conservation of metacorpus neurons provides high confidence in homology assignments between *H. glycines* neurons and their free-living counterparts (18).

In *C. elegans* and *P. pacificus*, the posterior esophagus forms a morphologically discrete terminal bulb containing seven esophageal neurons. In plant-parasitic nematodes, the region corresponding to the terminal bulb is dominated by three enlarged esophageal gland cells (**Fig. 1C and D**) (18). We identified seven neurons posterior of the isthmus in *H. glycines*. Unlike in *C. elegans* and *P. pacificus,* these posterior neurons are tightly clustered and differ in the anteroposterior position of individual neurons. Despite these differences, the conserved dorsoventral grouping of four dorsal and three ventral neurons, together with broadly similar neurite projection patterns (**Fig. 1C**; **Dataset S1**) support confident homology assignments to their free-living counterparts.

We observed one instance of intraspecific cell body position variation between the two *H. glycines* datasets. The M4 cell body is typically located in the metacorpus (8, 9, 20), but in one dataset (B3), it was displaced posteriorly into the isthmus region (**Fig. 1A**; **Fig. S2A**). Despite this variation, the anterior process of M4 consistently projected to the esophageal nerve ring in both individuals. This variation appears to be uncommon as the M4 cell body was consistently found in the metacorpus in two additional EM datasets, suggesting individual variability in neuronal cell body positioning in the *H. glycines* esophageal nervous system.

While the topological organization of neuronal cell bodies is conserved, most *H. glycines* esophageal neurons have relatively less complex branching patterns than their *C. elegans* homologs (**Fig. 1C**; **Dataset S1**). For example, the *H. glycines* I1 homolog lacks the anterior process present in both *C. elegans* and *P. pacificus*. In addition, the terminal bulb neurons of *H. glycines* lack all the posteriorly directed processes found in their *C. elegans* and *P. pacificus* homologs. M5 shows the most divergent morphology among the esophageal neurons across three species (**Fig. S2B**). In *C. elegans*, M5 innervates the grinder muscles and is restricted to the terminal bulb, whereas *H. glycines* and *P. pacificus* lack a grinder and their M5 homologs extend a long anterior process through the outermost dorsal nerve bundle along the length of isthmus.

Similar to *C. elegans* (20), we identified putative sensory endings adjacent to the esophageal lumen in *H. glycines*. While the relative positions of these endings were broadly conserved, we found only five endings from I3, M3L/R, and NSML/R in *H. glycines* (**Fig. S3**), whereas 15 of the 20 *C. elegans* neurons formed putative sensory endings (20). The reduced number of sensory endings may reflect a reorganization of esophageal sensory perception in *H. glycines*.

*H. glycines* exhibits two additional note-worthy modifications in the esophagus that are distinct from free-living species. First, in *C. elegans* and *P. pacificus*, the esophageal lumen lies on the central axis of the esophagus and runs straight along the entire esophagus. In contrast, the *H. glycines* procorpus lumen and adjacent neurons are displaced from the central axis and meander dorsoventrally along its length (**Fig. 1C and D**). This meandering course likely provides slack that allows the procorpus lumen to follow the stylet during stylet protraction. Second, the basement membrane and extracellular space separating the esophagus from other tissues is reduced in *H. glycines,* which may influence extra-synaptic communication between the esophageal and somatic nervous systems (**Fig. 1E**).

### Target organ connectivity is reduced and altered in *H. glycines*

While homologs of all 20 *C. elegans* esophageal neurons are present in *H. glycines*, the morphology of non-neuronal target tissues differ substantially. To test if these anatomical changes are associated with changes in synaptic connectivity, we identified and annotated chemical synapses and compared these to the established adult hermaphrodite *C. elegans* and *P. pacificus* esophageal connectome datasets (9, 20). To account for developmental differences between our *H. glycines* J2 data and the free-living adult datasets, we also reconstructed the esophageal connectome of a larval stage *C. elegans* molting into L2 from previously published EM data (19).

Consistent with previous findings in the somatic nervous system, the overall connectivity of the *C. elegans* esophagus is present in late L1, but with fewer connections than in the adult (**Fig. S4** and **Fig. S5**) (23).

*H. glycines* esophageal neurons innervate markedly fewer classes of cells than free-living species (**Table S1**). To compare the overall organization of synaptic output, we weighed each connection by its number of synapses and compared total synaptic output across target classes (**Fig. 2**A). In *H. glycines*, synapses targeting neurons (>60%) and glands (>12%) account for a larger fraction than in free-living species (<55% for neurons, approximately 6% for glands), whereas synapses targeting muscles comprise a smaller fraction (approximately 15%) than in free-living species (>31%). Marginal, epithelial cells and basement membrane targets contributed similar minor fractions in all species. Therefore, synaptic weight in *H. glycines* is shifted away from muscle toward neuronal and glandular targets, which aligns with its reduced musculature and enlarged gland cells.

**Fig. 2.**
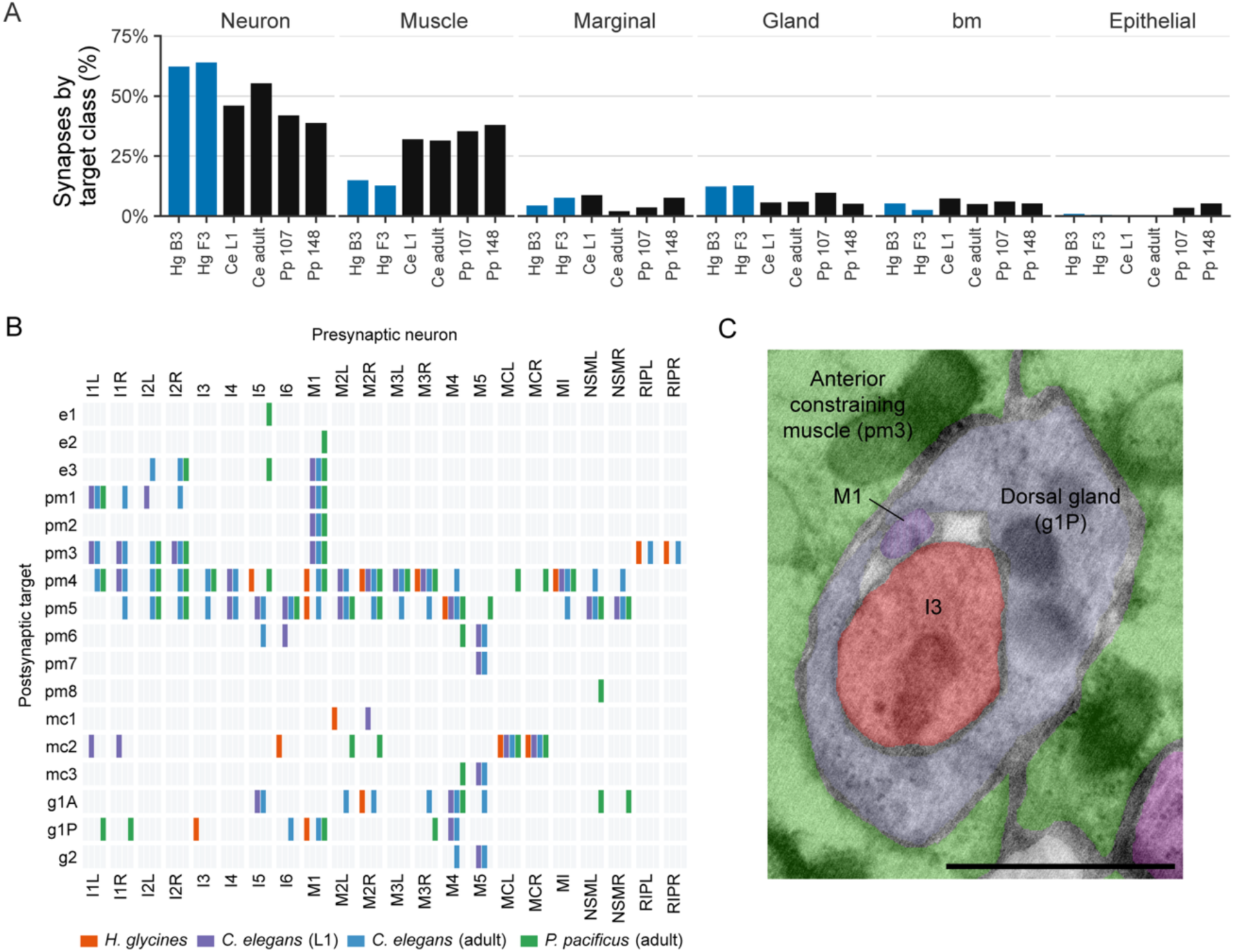
Comparison of synaptic output to target organs between *H. glycines* and free-living species (*C. elegans* and *P. pacificus*). (A) Proportion of weighted connections grouped by postsynaptic target classes. Connections were weighted by the number of synapses per connection. Blue, *H. glycines*; black, free-living species. Hg B3 and Hg F3, *H. glycines*; Ce L1, *C. elegans* late L1; Ce adult, *C. elegans* adult hermaphrodite, Pp 107 and Pp 148, *P. pacificus* adult hermaphrodite. (B) A connectivity matrix of directed chemical synaptic outputs to target organs. Columns represent presynaptic esophageal neurons and somatic RIP neurons, and rows represent postsynaptic target organ classes. Colored segments indicate the presence of a synaptic connection. Note that connections present in both individuals (shared connections) were used for *H. glycines* and *P. pacificus*. (C) Transverse EM image showing the dorsal gland process ensheathing M1 and I3. Scale bar, 1 µm.

We next examined the connectivity of individual target tissues associated with parasitic feeding behavior, including stylet-associated muscles, constraining muscles, and glands. To account for inter-individual variations, we restricted this comparison to connections present in both individuals for *H. glycines* and *P. pacificus* (see **Fig. S4** for complete connectivity matrix). The stylet- associated muscles (**Fig. 1A**), which drive stylet thrusting, are composed of two muscle classes, the stylet protractor and secondary stylet protractor muscles, proposed as homologous to e1 and e3 in *C. elegans* (**Fig. S2C**) (21). Unexpectedly, none of the esophageal neurons innervate the stylet protractor muscles in *H. glycines*, even though the homologous cells are innervated in *C. elegans* and *P. pacificus* (**Fig. 2B**). However, we observed a putative gap junction between RIP and the secondary protractor muscle (**Fig. S6**), although gap junctions were not systematically analyzed in this study.

*H. glycines* possesses two subventral glands and one dorsal gland homologous to *C. elegans* g1A and g1P (**Fig. S2C**), respectively. The *H. glycines* esophageal gland cells produce effector proteins essential for host penetration and feeding (24), while the esophageal gland cells of *C. elegans* have no apparent role in feeding (25). Synaptic input to gland cells shows considerable differences across species (**Fig. 2B**). For example, *H. glycines* has a unique I3 presynaptic connection to the dorsal gland (g1P). This connection is correlated with a unique ensheathment of I3 and M1 by the dorsal gland anterior process that is absent in free-living species (**Fig. 2C**). In *H. glycines,* the anterior process of the dorsal gland wraps the I3 and M1 processes along much of its length, which expands the local contact interface between the dorsal gland and these neurons.

*H. glycines* has two constraining muscle classes that lie immediately anterior and posterior of the metacorpus, homologous to *C. elegans* pm3 and pm5, respectively (**Fig. S2C**). Unlike these homologs, the *H. glycines* constraining muscles have myofilaments oriented tangential to the esophageal lumen and encircle gland processes, acting as a sphincter to regulate the transportation of gland cell vesicles during feeding (26). In *H. glycines*, the anterior constraining muscles are only innervated by the somatic neurons RIP, whereas the free-living species homologs show broader innervation from esophageal neurons (**Fig. 2B**). Synaptic inputs from RIP to pm3 were also found in adult *C. elegans* (20); however, our analysis suggests they are missing from the late *C. elegans* L1 stage. While we did not reconstruct the somatic nervous system connections to RIP in *H. glycines*, we located the cell body in a similar position to that in

*C. elegans* (**Fig. S7**). The absence of M1 innervation of pm3 in *H. glycines* is associated with reduced contact by dorsal gland ensheathment (**Fig. 2C**). M1 passes through the center of pm3 in *H. glycines*, but its anterior process is largely ensheathed by the dorsal gland extension, resulting in minimal contact with pm3. Like the anterior constraining muscles, the *H. glycines* posterior constraining muscles (pm5) receive synaptic input from a small subset of neurons (M1 and M4), whereas free-living species show broader innervation. Together, these data suggest a highly modified innervation in *H. glycines* compared to free-living nematodes.

### Extensive rewiring of the esophagus

To assess how the esophageal connectome has diverged, we quantified the similarity between datasets using the normalized edit distance (**Fig. 3A**). Within-species pairs were the most similar (0.19-0.42) across all pairwise comparisons, whereas pairs between *H. glycines* and free-living nematodes were more dissimilar (0.67-0.79), exceeding the distance between free-living species pairs (0.52-0.62). Because the connectomes in this study differ substantially in size (**Table S2**; **Fig. S8**), we compared each observed edit distance with those from degree-preserving random networks, which controls for size and degree. All within-species comparisons and comparisons across free-living nematodes were significantly more similar than the random expectation (adjusted P < 0.01; observed-expected, Δ = −0.113 to −0.472; **Fig. 3A**; **Table S3**). However, three of eight comparisons between *H. glycines* and free-living species did not differ significantly from random. The difference was larger within species (mean Δ = −0.357) than between free-living species (mean Δ = −0.117) or between *H. glycines* and free-living species (mean Δ = −0.084).

**Fig. 3.**
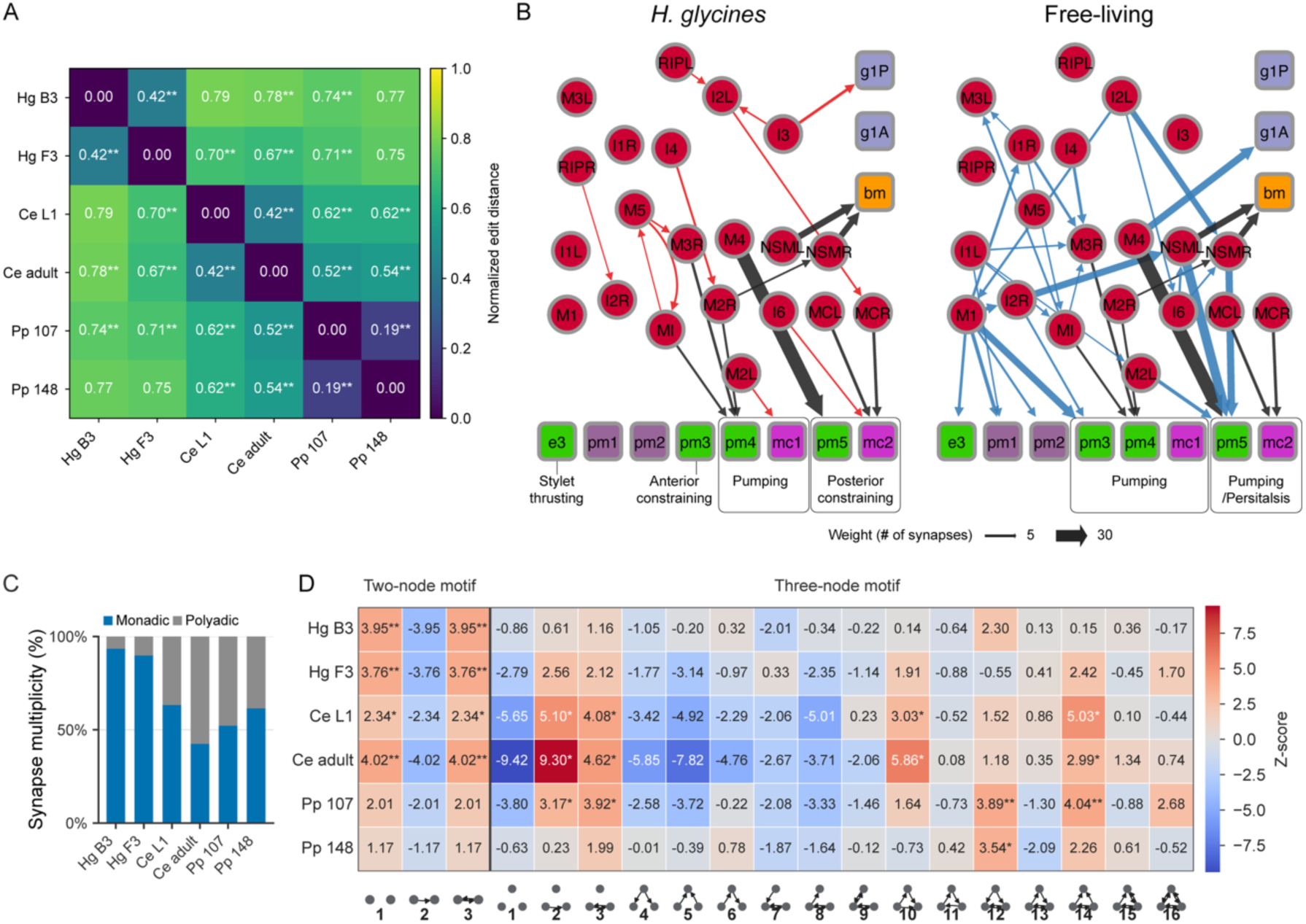
Divergence and rewiring of the esophageal connectome across nematode species. (A) Pairwise normalized edit distance matrix across six datasets (0 = identical connection sets, 1 = no shared connections). Observed normalized edit distances were compared to the distribution expected from degree-preserving randomized networks (n=500). **, p < 0.01. P-values were corrected using the step-down min-P algorithm. (B) Left, *H. glycines*-specific network (red, connections only present in both *H. glycines* individuals) and conserved network (black, connections present in all species). Right, free-living (*C. elegans* and *P. pacificus*)-specific network (blue, connections only present in free-living nematode datasets) and conserved network. Red nodes indicate neurons. Boxed nodes indicate non-neuronal targets: green, muscle; purple, epithelial cell; lavender, esophageal gland; magenta, marginal cell; yellow, basement membrane (bm). Non-neuronal targets are grouped by their associated function. e3, secondary stylet protractor; pm3, anterior constraining muscle; pm4, pump muscle; pm5, posterior constraining muscle; g1A, subventral gland; g1P, dorsal gland. Edge thickness indicates synaptic weight (averaged number of synapses per connection). (C) Proportion of monadic (blue) and polyadic synapses (gray). Polyadic synapses include a single synapse with more than one postsynaptic partner. (D) Heatmap of motif Z-scores for two-node motifs and three-node motifs across the six datasets. For each motif, the Z-score was computed against 1,000 degree-preserving randomized networks. Significance was determined by the step-down min-P procedure (*, p < 0.05; **, p < 0.01; ***, p < 0.001). Schematics below the columns depict the connectivity pattern of each motif. Hg, *H. glycines*; Ce, *C .elegans*; Pp, *P. pacificus*.

Thus, the divergence between *H. glycines* and free-living species greatly exceeds the variability between individuals of the same species.

The divergence prompted us to examine lifestyle-specific network subsets. We extracted connections present in both *H. glycines* datasets but absent from free-living datasets (*H. glycines*- specific, 11 connections), connections present in all free-living datasets but absent from *H. glycines* (free-living-specific, 27 connections), as well as connections present across all datasets (conserved, 9 connections) (**Fig. 3B**). Of the conserved connections, eight were concentrated in outputs to nonneuronal targets, including the pump muscle (pm4), posterior constraining muscle (pm5), marginal cells (mc2), and basement membrane. In contrast, both the *H. glycines*-specific and free-living-specific connections contained a higher proportion of neuron-neuron connections (63%-73%). The divergence is most apparent upstream of the pump muscle (pm4) output. In *H. glycines*, MI and M3R receive synaptic inputs from M5, and M2R from I4. In contrast, these neurons instead receive inputs from a different set of neurons in free-living specific subnetwork.

Therefore, while the connection from neurons to pump muscle is conserved, the upstream information flow into the conserved output module is divergent between *H. glycines* and free-living nematodes. By comparison, *H. glycines-*specific outputs to the gland and marginal cells receive little input within this subnetwork, indicating *H. glycines-*specific motor output rewiring without apparent changes in the upstream network.

### Network properties and predicted candidate neurons for *H. glycines* feeding control

#### H. glycines esophageal network is smaller than free-living species and monadic synapse-biased

Although the two *H. glycines* datasets differed in individual metrics, their network properties showed consistent differences from free-living species (**Table S2**). The *H. glycines* esophageal network contains fewer non-neuronal targets, fewer connections, and a lower average degree than both *C. elegans* late L1 and adult free-living species (**Fig. S9**). Consistent with having fewer targets, both the number of synapses and the synapse density are also lower in *H. glycines*. The most distinct difference in connectivity is a strong bias towards monadic synapses in *H. glycines* (**Fig. 3C**). In *H. glycines*, polyadic synapses (a single presynapse with multiple postsynaptic partners) account for approximately 10% or less, whereas free-living species have 37%-58% polyadic synapses. The prevalence of monadic synapses may allow each target class to be controlled independently in *H. glycines*.

Across all datasets, the entire esophageal network forms a single weakly connected component (every node is reachable from every other node in the undirected graph), whereas the largest strongly connected component (a group of nodes where every node is reachable in a directed graph) is smaller ranging from 8 to 18 nodes (**Table S2**). The small-worldness coefficient (*S*) ranged from 0.63 to 1.14 (27). *H. glycines* B3 (0.63) and F3 (0.99) did not exceed the threshold of 1, indicating it is not a small-world network. *H. glycines* has a longer average path length than free-living species and shows strong overrepresentation of reciprocity (2.5-3.2 fold relative to degree-preserving randomized networks), but lower transitivity and average clustering coefficient, which indicates a less integrated esophageal network in *H. glycines*.

#### Network motif analysis

To further investigate topological features, we compared the two-node and three-node motifs to degree-preserving randomized networks (28, 29). Motifs are recurring patterns of connectivity among small groups of nodes that represent the basic building blocks of a network. Statistical significance of motif overrepresentation varied across datasets in a species-specific manner (**Fig. 3D**). *H. glycines* and *C. elegans* showed significant overrepresentation of the reciprocal pair (motif 2). *H. glycines* showed no significant overrepresentation of any three-node motif, which is consistent with low clustering coefficient, whereas closed triangle motifs (motifs 10, 12, and 14) were overrepresented in free-living species. The overrepresentation of these triplet motifs is consistent with patterns reported in the *C. elegans* somatic nervous system and fruit fly brain network (29, 30). The enriched reciprocal pairs without overrepresented triplet motifs suggests that recurrent connectivity in *H. glycines* is concentrated in bidirectional neuron pairs rather than extending into three-node motifs. The feedback loop (motif 11) was not overrepresented in any dataset, which may indicate a relatively less complicated control structure in the esophagus.

#### Shifts in neuronal centrality and ablation predictions identify candidate neurons for H. glycines feeding behavior

To identify candidate neurons that may differentially influence feeding behavior in *H. glycines* and free-living species, we first compared centrality metrics (betweenness, PageRank, in-closeness, out-closeness) and in/out degrees in the esophageal network between *H. glycines* and free-living species (**Fig. 4A**; **Table S4**). Overall, *H. glycines* esophageal neurons showed lower degree than free-living species, resulting in reduced closeness. However, betweenness and PageRank increased in a subset of neurons, suggesting that information flow may be redistributed toward a smaller number of neurons, despite the overall loss of connectivity. M5 was the only neuron whose mean value was higher in *H. glycines* across all metrics and degrees. Consistent with loss of non-neuronal targets, the *H. glycines* I2 neuron had reduced out-degree but increased mean PageRank compared to free-living species. In *H. glycines*, I2 receives chemical synapses from RIP that may replace the RIP-I1 gap junction in free-living species. Conversely, I1L and M3L/R were reduced across all centrality metrics in *H. glycines*. I1 lacks the anterior process and has lost its connection with RIP in *H. glycines*. M3 is a motor neuron that synapses only onto pm4 in *H. glycines*. Thus, centrality shifts reflect both a global reduction of the esophageal network and neuron-specific changes in morphology and connectivity.

**Fig. 4.**
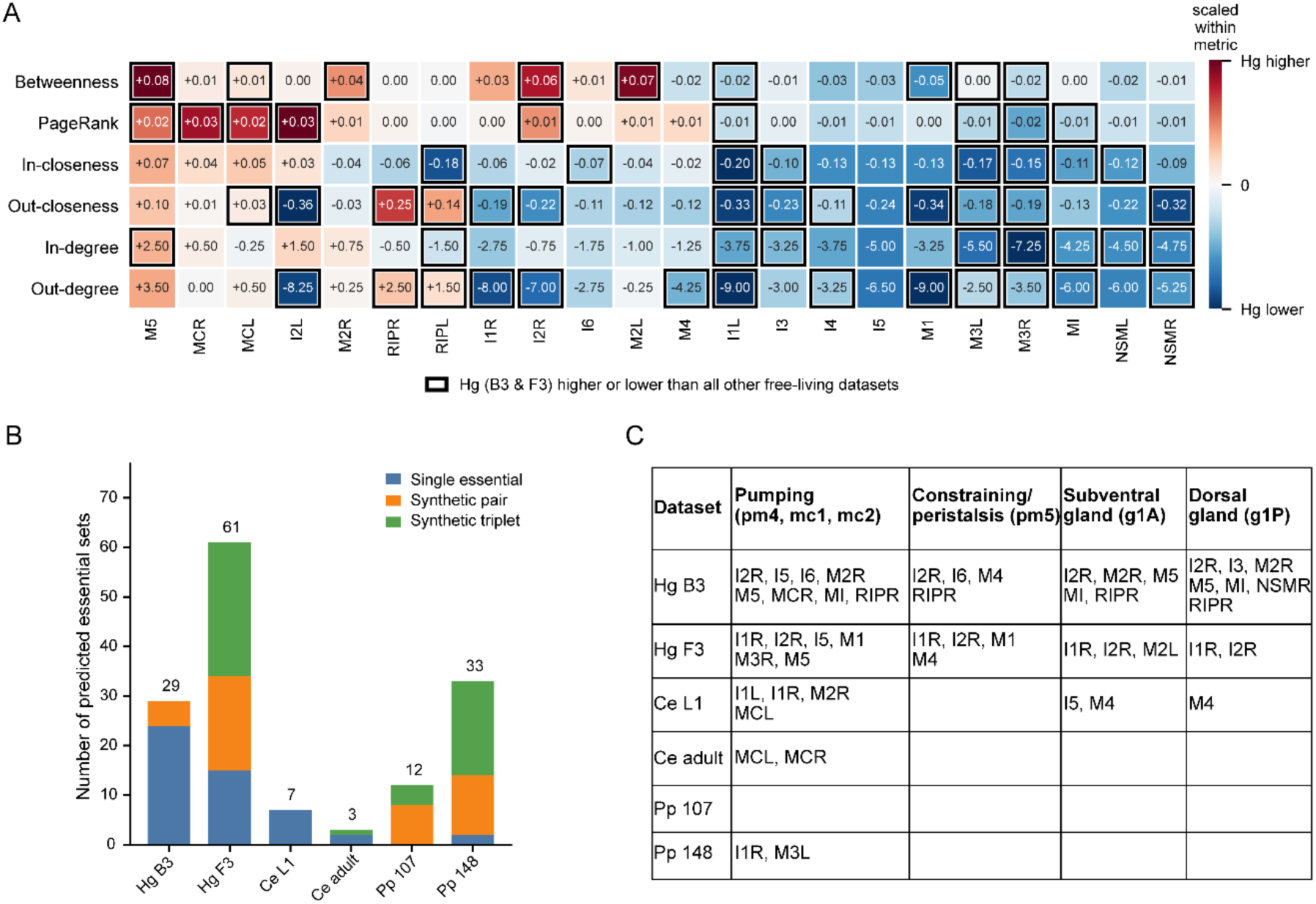
Comparison of centrality metrics and degrees between *H.* glycines and free-living species (*C. elegans* and *P. pacificus* adult) and prediction of essential neurons by synthetic ablation analysis. (A) Cell values represent the mean differences in metric between *H. glycines* (mean of B3 and F3) and free-living species (mean of free-living datasets). Red cells indicate higher centrality in *H. glycines*; blue cells indicate lower centrality. Raw differences were scaled within each metric by dividing by the maximum absolute raw difference in the metric, giving values from -1 to +1. Black-bordered cells indicate consistent differences between *H. glycines* and free-living species across all six datasets (see **Table S4**). (B) Number of predicted essential sets identified by synthetic ablation analysis. Sensory input was assumed to enter through RIPL/R in *H. glycines* and through I1L/R free-living species. (C) Predicted single essential neurons for each of the four target modules, including pumping (pm4, mc1, mc2), constraining/peristalsis (pm5), subventral gland (g1A), and dorsal gland (g1P). Hg B3 and Hg F3, *H. glycines*; Ce L1, *C. elegans* late L1; Ce adult, *C. elegans* adult hermaphrodite, Pp 107 and Pp 148, *P. pacificus* adult hermaphrodite.

To further identify neurons potentially involved in specific feeding circuits, we performed synthetic ablation analysis (31). This analysis predicts controllability (see SI Appendix, Extended Materials and Methods) to target nodes to identify single essential neurons whose removal reduces the output score, as well as synthetic essential pairs and triplets that are individually non-essential but become essential when removed in combination. We first confirmed that the analysis results in known essential neurons in *C. elegans*. In *C. elegans* adult, we identified MCL/R as essential neurons for the pumping module, which agrees with the pumping defect of MC-ablated animals (**Fig. 4C**) (32). In *H. glycines*, multiple single, pair, and triplet combinations were detected, and more single essential neurons were found across all modules than in the free-living datasets (**Fig. 4B**; **Table S5**). While the number of predicted combinations differed between the two *H. glycines* individuals, I2R was predicted to be essential across all four target modules in both specimens.

Similarly, M5 and I5 were predicted as essential neurons in the pumping module, and M4 in the constraining module in both individuals. In contrast, neurons with reduced centrality, such as I1L and M3L, did not show consistent intra- or interspecific patterns.

## Discussion

### Simplification and rewiring of the *H. glycines* esophageal connectome

Despite an estimated divergence time of more than 300 million years (11) and clear behavioral and morphological differences between *H. glycines* and free-living nematode species, we found remarkable conservation of the esophageal nervous system organization. Properties from cellular composition, including the number of neurons, neuronal identity, neurite projection pattern to overrepresentation of reciprocal motifs are all maintained across species, which suggests principles that are broadly conserved across the nematode esophageal nervous system. Against this conservation, however, we reveal parasite-adapted neural circuits. Our findings demonstrate several instances of evolutionary modifications on a conserved anatomical scaffold in *H. glycines* J2. We find that *H. glycines* has reduced chemical synapses and rewired conserved structures through selective partner reassignment.

Nervous system simplification is common among diverse animal parasites. Parasitic flatworms appear to have reduced neural complexity in the peripheral nervous system relative to their free-living relatives (2). The parasitic wasp *Megaphragma* undergoes neuronal loss during development, resulting in 95% anucleate neurons in the adult stage (3). Most dramatic is the complete absence of a nervous system in Myxozoa cnidarians (1, 33). Our data suggests this simplification during the evolutionary transition from free-living to parasite can extend to chemical synaptic networks, even when the total number of neurons is equivalent. In *H. glycines,* the reduction in connectivity is most prominent in the neuron-muscle connection. Interestingly, neuron-muscle connections present in both *H. glycines* datasets are also shared with at least one other species. This suggests that neuron-muscle connectivity likely evolved through the elimination of existing synapses, rather than through gain of novel connections.

In contrast to the reduction of synaptic connectivity, the weighted increase of connections to the gland cells corresponds with the increased importance of these cells during infection. Recent work has demonstrated that a master transcriptional regulator of gland cell effectors is directly activated by sensing plant-derived signals (34), suggesting a functional connection between sensory input and gland cell function during parasitism. Modifications to neuronal input of gland cells is also evidenced through the ensheathment of neurons by the dorsal gland process. This ensheathment is reminiscent of glial wrapping in several species. In *C. elegans*, the amphid sheath cells are glia with known secretory function that wrap the distal ends of sensory neuron dendritic processes (35). Unlike the amphid sheath cells, ensheathment of I3 and M1 by the dorsal gland process occurs along the entire anterior half of its process. This glial-like ensheathment was also observed in a different plant-parasitic nematode, *Meloidogyne incognita* (17), suggesting that the gland cell ensheathment of neurons is an evolutionary modification relevant to plant parasitism.

### The esophageal connectome of plant-parasitic nematodes provides insights into the development of new targeted control strategies

Comparative analysis of the esophageal connectome can identify parasitism-specific neuronal adaptations that can be selectively targeted for control of plant-parasitic nematodes. Targeting the signaling pathways mediating *H. glycines-*specific connections (e.g., dorsal gland-I3, mc1-M2L, I6-mc2) is expected to yield effects that are selective for *H. glycines*. In particular, I3 forms a putative sensory ending near the dorsal gland ampulla, potentially contributing to regulation of gland secretion. Beyond direct connections to motor/gland outputs, neurons predicted to be associated with the reorganization of network information flow can be alternative candidates. Centrality shifts, morphological divergence, changes in target connectivity, and synthetic ablation predictions (31) all converge on I2R and M5 as primary candidate neurons in *H. glycines*. These findings suggest that the *H. glycines* esophageal network has been reorganized such that feeding control is concentrated around a smaller number of neurons.

Our reconstruction will provide an essential cellular map for future molecular localization studies. While neuronal identity is conserved across nematode species, the neurotransmitter receptor profile is highly divergent even within a single genus (36). Use of our anatomical map combined with analysis of molecular receptors may lead to the identification of highly specific potential candidates for control. In *C. elegans*, serotonin stimulates increased esophageal pumping (37). In plant-parasitic nematodes, serotonin instead stimulates stylet thrusting, but not corresponding pumping (38–40). Because neuronal identity is broadly conserved but serotonin drives distinct feeding behaviors, we speculate that the serotonergic pathway may have been functionally reassigned through divergence in serotonin receptor profiles.

To integrate these findings with the development of future control strategies, further research is needed to validate individual neuronal functions. Although the esophageal connectome identified in this study is limited to *H. glycines*, esophageal morphology is very similar among the economically important Tylenchomorpha species of plant-parasitic nematodes (17), suggesting that these findings may be extended beyond this species. However, it is important to note that our comparison is limited to three species within the subclass Chromadoria. The phylum Nematoda encompasses a wide range of morphological and ecological diversity and plant parasitism has evolved independently several times (13). Recently, we reported substantial increase in the number of neurons in the ventral nerve cord and the body wall sensory system in the early-diverging subclass Dorylaimia, compared to more recently derived lineages like *C. elegans* (41). To determine whether the simplification of the esophageal connectome observed in this study is a general consequence of the shift to a plant-parasitic lifestyle or a phenomenon specific to the *H. glycines* lineage, additional connectome data from other lineages is required.

Unlike *C. elegans* and *P. pacificus, H. glycines* only reproduces sexually. While our *H. glycines* strain has been maintained as a highly inbred greenhouse line for over a decade, substantial genetic diversity within populations is likely (42). The two individuals we reconstructed exhibited considerable differences in synaptic connectivity (**Fig. S1**). Despite these differences, the overall connectivity patterns and network metrics were similar. Witvliet et al. (23) reported that approximately 43% of connections were not conserved between isogenic individuals of *C. elegans*. In our data, we found a similar proportion of variable connections between individuals. We therefore infer that the synaptic variability attributable to genetic variability is unlikely to exceed the stochastic variability reported previously.

This study focused solely on chemical synapses. In the *C. elegans* esophagus, gap junctions between adjacent muscle cells play a crucial role in the rapid propagation of contraction signals (43). The lack of chemical synaptic input to the stylet-associated muscles suggests that these muscles are likely electrically coupled by gap junctions (**Fig. S6**) or mediated by neuropeptide. However, identification of gap junctions from EM is extremely challenging due to their often variable and ambiguous morphology, thus often excluded from analysis for comparative connectomics (4, 6, 23). Instead, chemical synapse identification is relatively stable and offers reproducible data.

## Materials and Methods

Detailed materials and methods are provided in SI Appendix, Extended Materials and Methods.

*H. glycines*, originally isolated from Illinois, was maintained on soybean (*Glycine max* Williams 82) in the greenhouse. Eggs were extracted by sucrose-centrifugation method (44) and incubated in hatching chambers for 24 hours to collect freshly hatched J2s. We generated four serial-section EM datasets of *H. glycines* J2 and used two of them to reconstruct the esophageal connectome of two individuals (**Fig. S1**). Both are transverse datasets, covering the esophagus, comprising 1,620 sections (dataset F3) and 1,775 sections (dataset B3) and spanning from the head of the nematode to the posterior end of the esophageal gland cells. The remaining two datasets, one transverse dataset partially covering the region from the posterior procorpus to the anterior isthmus, and one longitudinal dataset covering the entire isthmus were used to confirm observations from datasets B3 and F3.

## Acknowledgments and funding sources

We thank Jonathan Boudreaux, Hailey Liu, Dzhuliia Vago, Darian Figueroa, Bridget Neira, and Matt Barnum for their assistance in EM image acquisition and segmentation. We also thank the University of Illinois Beckman Institute Imaging Technology Group staff Cate Wallace and T. Josek for their assistance in utilizing HPF and EM. Funding was from the USDA-NIFA 2021- 67013-33737.

## Author Contributions

L.R.V. and N.E.S. designed research; J.H., A.R.T., and E.C. performed research; J.H., A.R.T., E.C., L.R.V., and N.E.S. analyzed data; J.H. and N.E.S. wrote the paper.

## Classification

Biological Sciences; Neuroscience

## Extended Materials and Methods

### Serial-section scanning EM

High-pressure freezing (HPF) was performed according to Hall et al. (1) with minor modifications.

The freshly hatched J2s were placed in a 100-µm deep HPF planchette coated with 1- hexadecene and filled with 20% bovine serum albumin, then frozen using a Bal-tec HPM 010 high-pressure freezer. Subsequently, the specimens were moved to 2% OsO4, 0.1% uranyl acetate, and 2% H_2_O in acetone in an FS-8500 freeze substitution unit. The temperature program was as follows: -90 °C for 110 hours, ramped at 4 °C/hour to -20 °C for 16 hours, then ramped at 4 °C/hour to 0°C. Samples were washed with pre-chilled 100% acetone for 15 minutes for three times at 0 °C and three additional times at room temperature. Samples were transferred to microporous capsules and infiltrated with Polybed812 resin (Electron Microscopy Sciences) in a stepwise series: 1:1 resin:acetone for 24 hours, 2:1 resin:acetone for 24 hours, and 100% resin for 24 hours on an orbital shaker at room temperature, followed by three changes of 100% resin over 8 hours. Samples were then embedded in resin-filled molds or flat embedded within a resin-filled Gene Frame (ThermoFisher) between Aclar (Ted Pella) films (2) and polymerized at 60°C for 48 hours.

Serial 70-nm sections were cut using a PowerTome PC ultramicrotome with an Ultra ATS diamond knife (DiATOME) and collected on silicon wafers (Ted Pella). Silicon wafers were cut into 2x2 cm rectangles and glow discharged for 1 min before use (Denton Vacuum). Serial sections were post-stained with 2% uranyl acetate for 30 minutes followed by Reynold’s lead citrate for 5 minutes. The esophageal region of the sections was then imaged using a FEI Quanta FEG 450 ESEM with a backscatter detector. Images were acquired at 3.37-nm/pixel (10,000-20,000x magnification).

### Reconstruction of esophageal connectome and chemical synapse annotation

Raw images were montaged and aligned using TrakEM2 (3) and imported into VAST Lite (4) for annotation and segmentation. All esophageal neurons were volumetrically segmented and annotated in both *H. glycines* F3 and B3 datasets. Presynaptic active zones were segmented for chemical synapse annotation. To ensure reliability of synapse annotations, two annotators independently performed synapse annotations. The annotations were then cross-checked by both annotators and any discrepancies were resolved by a third annotator. Only synapses agreed upon by all three were used as the final synapse connectivity dataset.

Previously established *C. elegans* late L1 EM dataset was acquired from https://doi.org/10.60533/BOSS-2022-1JI1 (5) and imported into VAST Lite for volumetric reconstruction and chemical synapse annotation following the same procedures described above for *H. glycines*. The volumetric reconstruction of the *C. elegans* late L1 esophagus is available at WormFindr (https://wormfindr.web.illinois.edu/v1/). Chemical synaptic connectivity data for adult hermaphrodite *C. elegans* and *P. pacificus* were obtained from previously published sources (6, 7).

### Network construction

> Each connectome was represented as a directed graph in which nodes are neurons and non-neuronal target cells, and directed edges are chemical synaptic connections from presynaptic to the postsynaptic cell. Polyadic synapses (a single presynapse with multiple postsynaptic partners) were decomposed into separate directed pre-postsynaptic pairs, one for each postsynaptic partner, with each pair contributing one unit of synaptic weight. Unless stated otherwise, every analysis was performed on each dataset individually. Individual connectivity matrices, wiring diagrams, and target tables are provided in **Fig. S4; Fig. S5**; **Table S1**.

### Target organ connectivity

For reliable and conservative cross-species comparison of target organ connectivity, we constructed shared network for *H. glycines* by extracting the connections present in both individual datasets B3 and F3. For *P. pacificus,* a shared network reconstructed from specimens 107 and 148 was used (6). The *C. elegans* L1 network derives from a single individual and the adult dataset is an established composite network (7). Because some non-neuronal targets are subdivided into radial sectors that differ in number or relative radial position among species, radial sectors of the same target cell were collapsed into a single node.

To compare the distribution of synaptic output among postsynaptic target classes, each connection was assigned to the cell type of its postsynaptic cell (neuron, muscle, marginal cell, epithelial cell, gland, or basement membrane). For each dataset, we then computed the proportion of connections in each type as an unweighted fraction of all connections and as a fraction weighted by synaptic weight.

### Edit distance

Pairwise dissimilarity between connectomes was quantified as a normalized edit distance on the binary directed networks. For two edge sets *E*_1_ and *E*_2_, the normalized distance was |*E*_1_Δ*E*_2_|/(|*E*_1_| + |*E*_2_|). Radial sectors of the same target cell were collapsed into a single node. The observed edit distances were tested against a degree-preserving null model. For each pair of networks, we generated 500 randomized networks and recomputed the edit distance for each randomized pair. Each observed distance was expressed as a Z-score relative to the null distribution. P-values were corrected using a step-down min-P procedure.

### Lifestyle-specific connection subsets

A connection was classified as *H. glycines*-specific if present in both *H. glycines* individuals but absent from all free-living datasets. Free-living-specific connection are those present in all free-living datasets but absent from *H. glycines*. A connection was classified as conserved if present across all datasets.

### Network metrics

Network density was E/[N(N-1)], where E is an edge and N is a node. Reciprocity was computed as the fraction of edges with a reciprocal counterpart. Transitivity was the fraction of all possible triangles in the graph. The average clustering coefficient is the probability that, for three neurons, given that one neuron is connected to two others, those two are also connected to each other.

The average path length is the mean shortest path length over all node pairs, and the diameter is the longest of those shortest paths on the undirected network. Reciprocity, transitivity, average clustering coefficient, and average path length were compared with averages over 500 degree-preserving random graphs.

We report the sizes of the largest strongly and weakly connected components (SCC and WCC) and within the largest SCC, the mean directed shortest path length and average clustering coefficient. The small-worldness coefficient (*S*) was calculated as 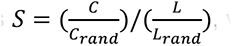, where C and L are the clustering coefficient and characteristic path length of the largest SCC, and C_rand and L_rand are their averages over 500 directed Erdős–Rényi random graphs (8).

Each synapse was classified by the number of postsynaptic partners as monadic (one partner) or polyadic (two or more). A connection was defined as a unique pre- and postsynaptic pair.

Synaptic weight was defined as the number of synapses assigned to a pre- and postsynaptic cell pair. For polyadic synapses, each postsynaptic partner received equal synaptic weight of 1. Total synaptic weight is the total number of synapses across all connections.

### Degree and connection weight distributions

Edges sharing the same presynaptic cell and the same postsynaptic class were merged into a single directed edge (connection), and their synapse counts were summed. In- and out-degrees were defined as the number of unique incoming and outgoing edges of a node, respectively, and total degree was the sum of the two. Degree distributions included all neuronal and non-neuronal nodes. Empirical complementary cumulative distribution functions (cCDF) were calculated as the proportion of observations exceeding each integer threshold *k*, P(X>*k*). The two *H. glycines* datasets were compared using two-sample Kolmogorov-Smirnov (KS) tests. For comparisons among the six datasets, overall differences were assessed using Kruskal-Wallis tests, followed by all pairwise two-sided Mann-Whitney U tests and two-sample KS tests. P values were adjusted using the Holm procedure.

### Centrality analysis

Unweighted four centrality measures (betweenness, PageRank, in-closeness, and out-closeness) were on the directed esophageal network. Radial sectors of the same non-neuronal target cell were collapsed into a single node. In- or out-degree was defined as the number of presynaptic neuron or postsynaptic neuron, respectively. For comparison, each measure was averaged within *H. glycines* (B3 and F3) and within the free-living datasets, the difference (*H. glycines* – free-living) was used for generating the heatmap. The mean difference above or below 0 indicates increased or decreased centrality in *H. glycines*, respectively. Raw differences were scaled within each metric by dividing by the maximum absolute difference in the metric, giving values from -1 to +1.

### Motif analysis

Two-node and three-node motif analyses were performed on networks including both neuronal and non-neuronal nodes. Observed motif counts were compared against a null distribution derived from 1,000 degree-preserving randomized networks (9). For each motif class, one-sided upper tail P values were calculated by comparing the observed motif count with the corresponding null distribution from randomized networks and corrected for multiple comparisons using the step-down min-P procedure (9). Motif counts were converted to z-scores by comparing the observed count for each motif with the corresponding null distribution from randomized networks.

### Synthetic ablation analysis

Synthetic ablation analysis was performed using a target-control maximum-flow framework (10, 11). For each dataset, directed chemical synapse graphs were converted to binary connectivity graphs. The input was set to RIPL/R for *H. glycines* J2 and I1L/R for *C. elegans* and *P. pacificus*. While RIP is not present as an input in the chemical esophageal network in free-living species, RIP is coupled with I1 by gap junctions (6, 12). Four output modules were analyzed separately and comprised pumping (pm4, mc1, mc2), constraining/peristalsis (pm5), subventral gland (g1A), and dorsal gland (g1P). Because pm4 and pm5 are single syncytia in *C. elegans* and *P. pacificus*, radial sectors of these muscles were grouped into each muscle class.

For each output module, analysis was restricted to the directed subnetwork defined as the intersection of nodes reachable from the input set and nodes that could reach the output set. A time-expanded linking graph was then constructed in which each node was split into unit-capacity copies across temporal layers. Input neurons were connected to a super-source, output nodes were connected to a super sink, and the maximum flow from source to sink was used as the baseline controllability score, *R*_0_. Neuronal ablation was modeled by deleting all temporal copies of the ablated neuron or neuron set and recalculating the maximum-flow score, *R*(*A*).

A single neuron was classified as essential if its removal reduced the score below *R*_0_. A synthetic essential pair was defined as two individually nonessential neurons whose combined removal reduced the score. A strict synthetic essential triplet was defined as three individually nonessential neurons for which no pair subset reduced the score, but simultaneous ablation of all three neurons reduced the score. Combinations that removed the entire input set were excluded from synthetic essential counts.

### Software and code availability

Raw EM images were montaged and aligned using TrakEM2 (3). Neurons and synapses were segmented in VAST Lite (4). Data analyses were performed in Python 3.13.9 using NetworkX 3.5, NumPy 2.3.5, pandas 2.3.3, Matplotlib 3.10.6, seaborn 0.13.2, SciPy 1.16.3, statsmodels 0.14.5, openpyxl 3.1.5, joblib 1.5.2. Wiring diagrams were generated using Cytoscape 3.10.3. Three-dimensional reconstructions were rendered in Blender 4.5. The raw EM image volumes for the *H. glycines* B3 and F3 datasets are currently being deposited in BossDB.org and will be made publicly available upon completion of repository ingestion. All scripts and files used to generate figures are available at https://github.com/jhan011/hglycines-esophageal-connectome.

**Fig. S1.**
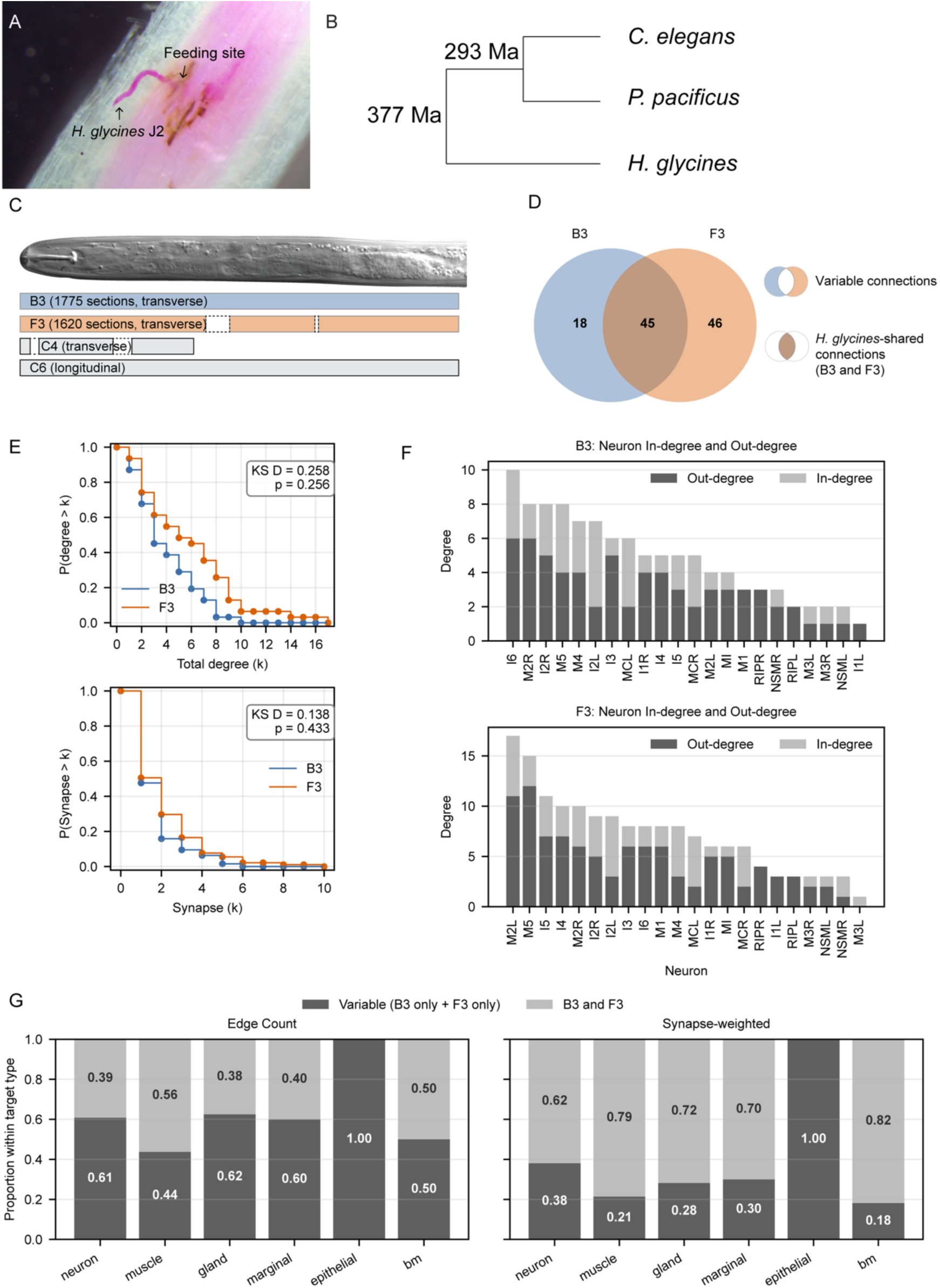
Comparison of *H. glycines* esophageal connectome datasets. (A) Light micrograph of a J2 *H. glycines* feeding within a soybean root. The *H. glycines* was stained in red with acid fuchsin and the root cortex cleared with bleach. (B) Phylogeny of *C. elegans, H. glycines,* and *P. pacificus* with estimated divergence dates in million years (Ma) (13). Branch lengths do not represent phylogenetic distance. (C) DIC micrograph of a J2 *H. glycines* with approximate coverage of four serial-section EM datasets. B3 and F3 were reconstructed for analysis. C4 and C6 were used for additional anatomical reference. Dashed lines indicate gaps in the dataset. (D) Variable (connections present only in B3 or F3) and shared (B3 + F3) connections (unique pre- and postsynaptic pair). Edges sharing the same non-neuronal postsynaptic class were collapsed into a single connection. (E) Complementary cumulative distributions of degree (top) and the number of synapses (bottom) in *H. glycines* B3 and F3. Distributions were compared by a two-sample Kolmogorov-Smirnov test. (F) In-degree and out-degree distributions of individual esophageal neuron and RIPL/R. (G) Proportion of variable and shared connections (left), synapse-weighted connections (right). Weighted shared connections account for a larger fraction than unweighted connection count. Bm, basement membrane.

**Fig. S2.**
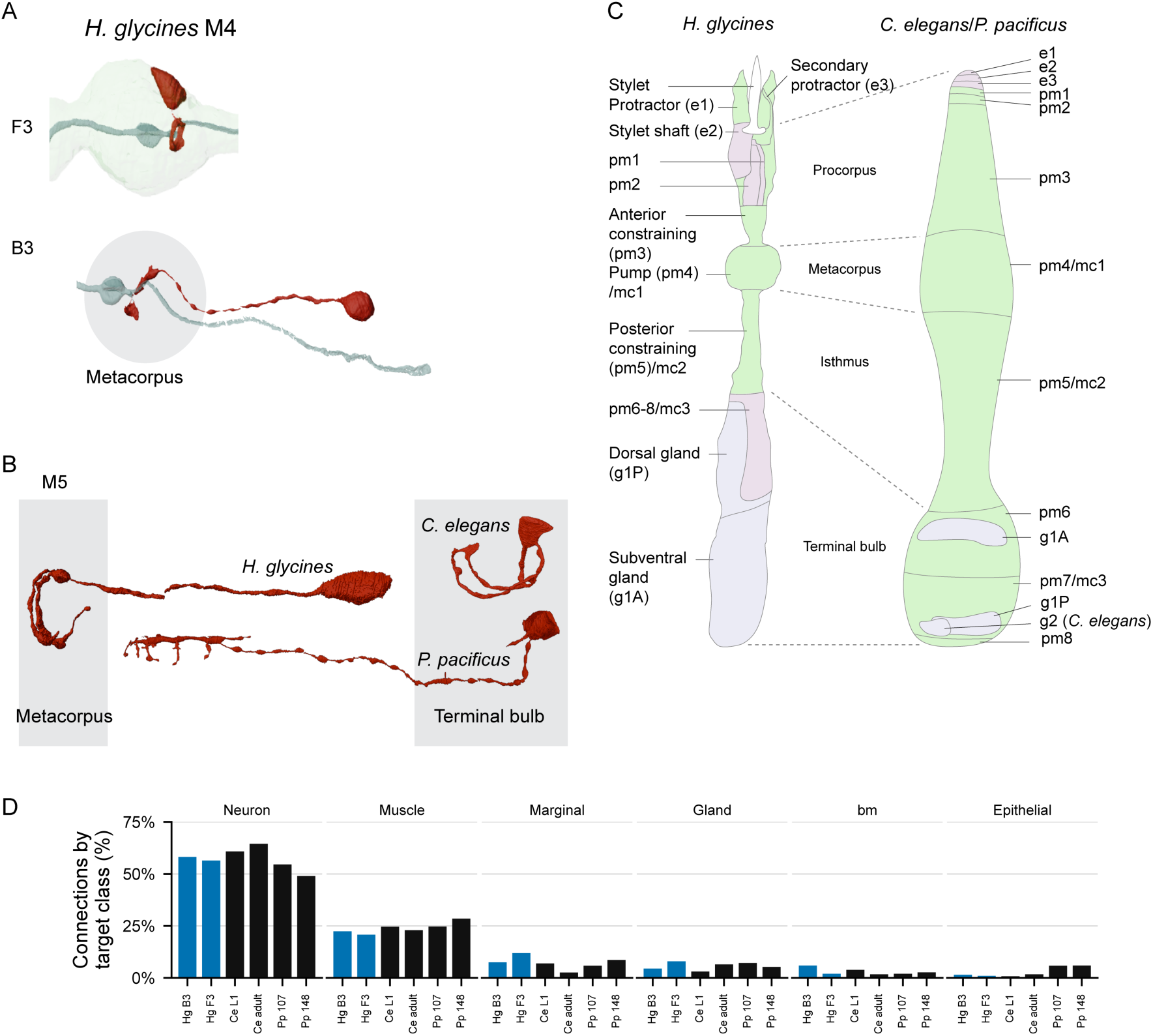
Variation in esophageal anatomy and target connectivity proportion. (A) Variation in the position of M4 between different specimens of *H. glycines* J2. The process remains within the metacorpus in both individuals. (B) Morphological differences of M5 across *H. glycines* J2, *C. elegans* late L1, and *P. pacificus* adult hermaphrodite. Gray boxes indicate the approximate positions relative to the metacorpus and terminal bulb. Note that the scale differs among species, so sizes are not directly comparable. The *C. elegans* and *P. pacificus* reconstructions were generated from previously published EM data (5, 6). (C) Simplified schematic diagrams of the esophagus in *H. glycines* (left) and *C. elegans* and *P. pacificus* (right). Green, muscle; purple, epithelial cell; lavender, gland cell; bm, basement membrane. (D) Proportion of connections (unique presynaptic and postsynaptic pairs) grouped by postsynaptic target classes. Blue, *H. glycines*; black, free-living nematodes.

**Fig. S3.**
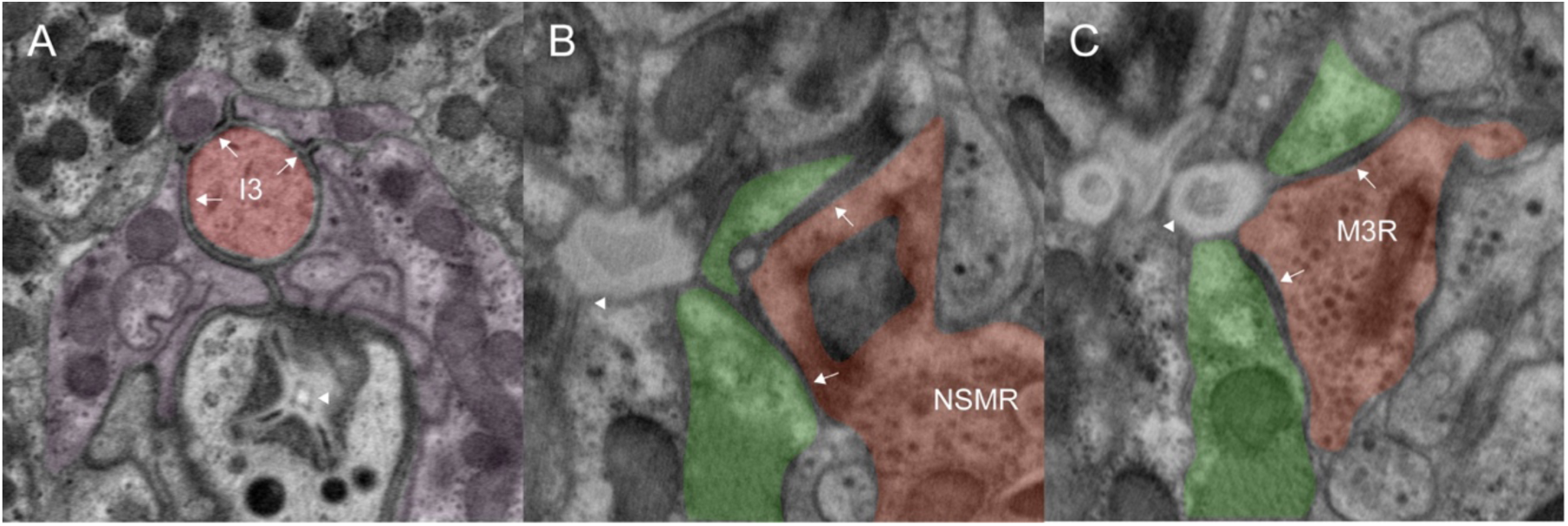
Putative sensory endings of *H. glycines* esophageal neurons. (A) The apical anterior tip of I3 forming adherens junctions (AJ) with pm1 and pm2 near the dorsal gland ampulla valve (arrowhead). (B) NSMR forming AJs with pm4 and pm5. Arrowhead: lumen. (C) M3R forming AJs with pm4 and pm5. Arrowhead: lumen branch immediately before the subventral gland ampulla valve. Arrows indicate AJs. Red: neuron; purple: epithelial cell; green: muscle.

**Fig. S4.**
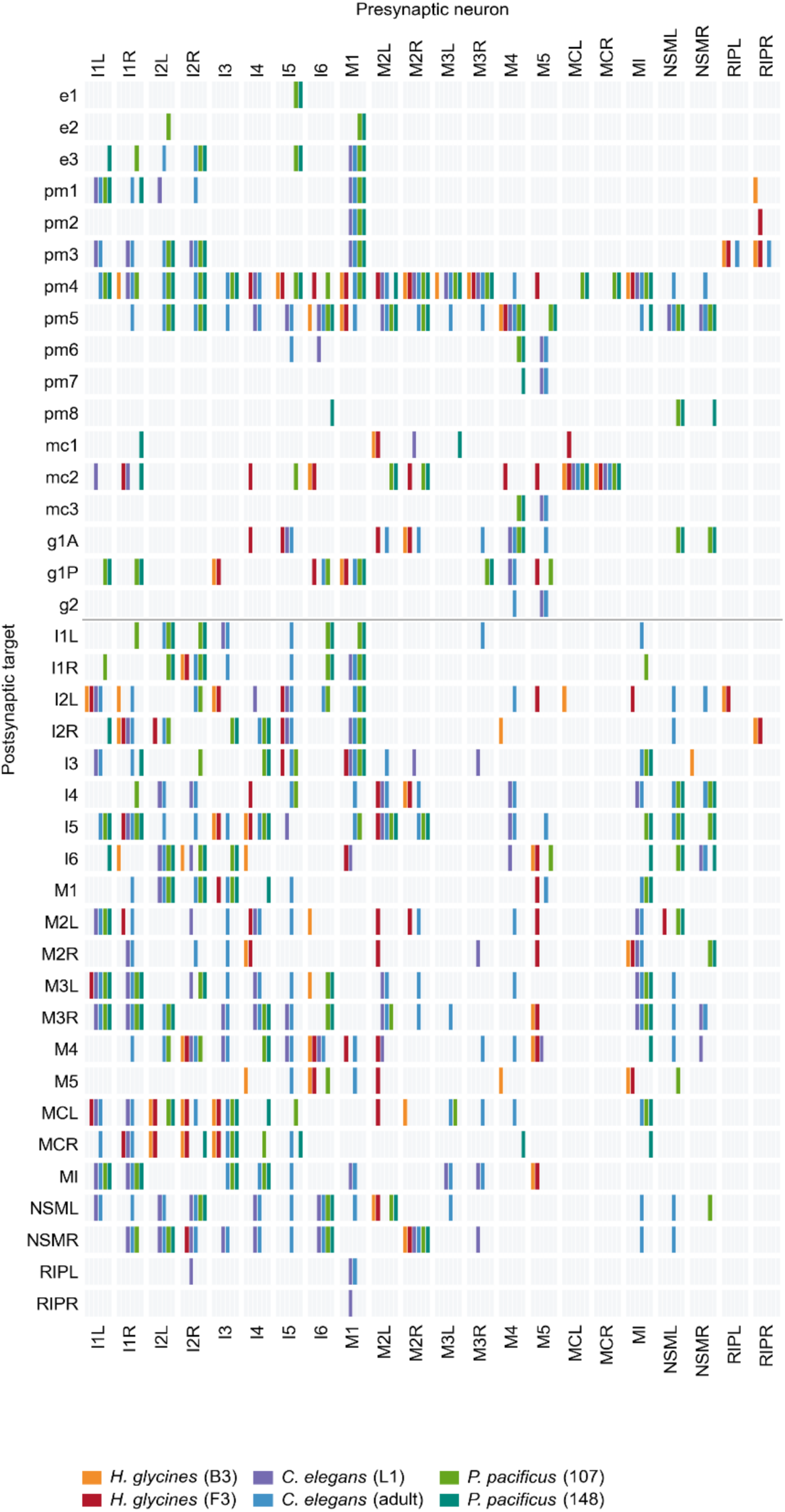
Connectivity matrix of the chemical synapse esophageal networks. Columns represent presynaptic esophageal and somatic RIP neurons, and rows represent postsynaptic partners. Non-neuronal targets are grouped by class. Colored segments indicate the presence of a synaptic connection.

**Fig. S5.**
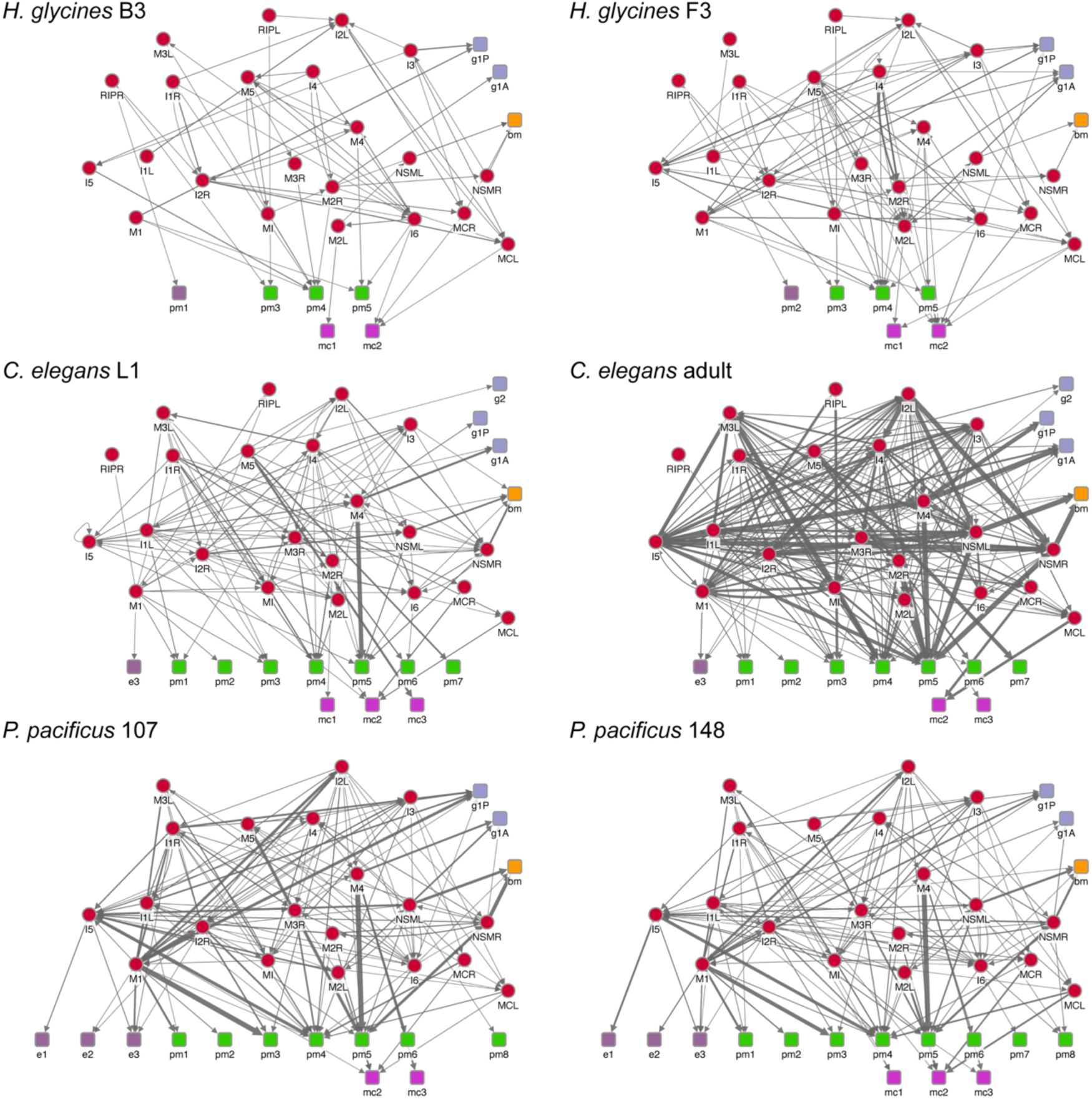
Wiring diagrams of the chemical esophageal network of individual datasets. Red, neuron; green, muscle; purple, epithelial cell; lavender, gland cell; yellow, basement membrane (bm). Descriptions in *H. glycines*: e1, stylet protractor muscle; e3, secondary stylet protractor; pm3, anterior constraining muscle; pm4, pump muscle; pm5, posterior constraining muscle; g1A, subventral gland; g1P, dorsal gland. Edge thickness indicates synaptic weight (number of synapses).

**Fig. S6.**
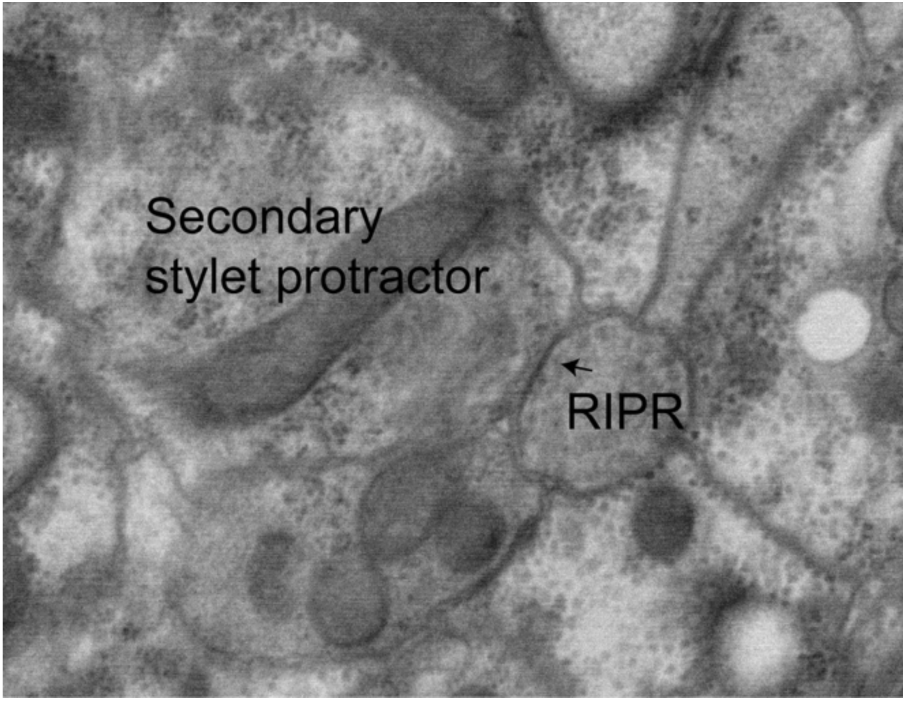
Transverse EM image of a putative gap junction (arrow) between RIPR and the secondary stylet protractor muscle (*C. elegans* e3 homolog) in *H. glycines*.

**Fig. S7.**
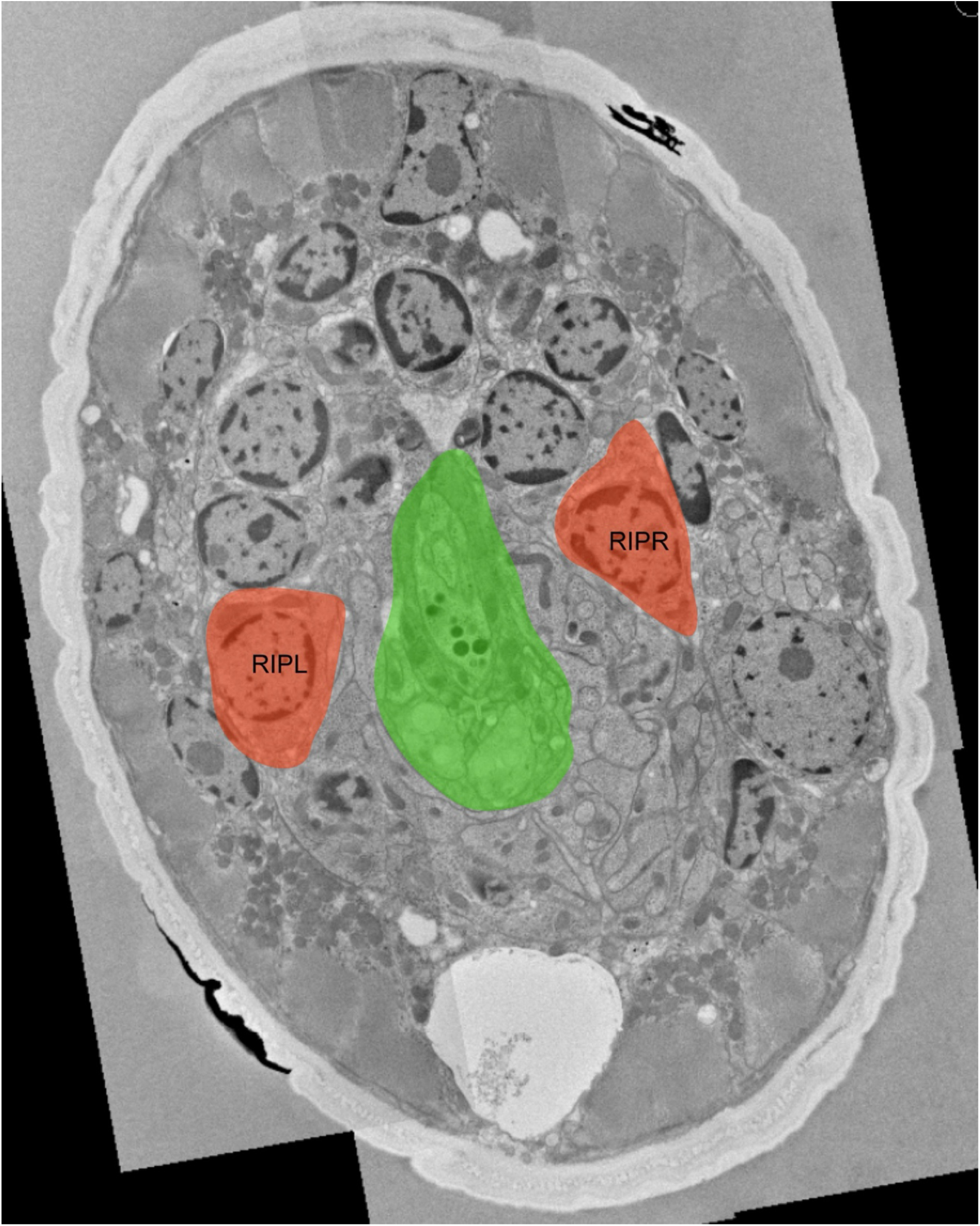
Transverse EM image of *H. glycines* J2 showing the esophagus (green) and cell bodies of putative RIP homologs (red) in the subdorsal sectors at the anterior isthmus.

**Fig. S8.**
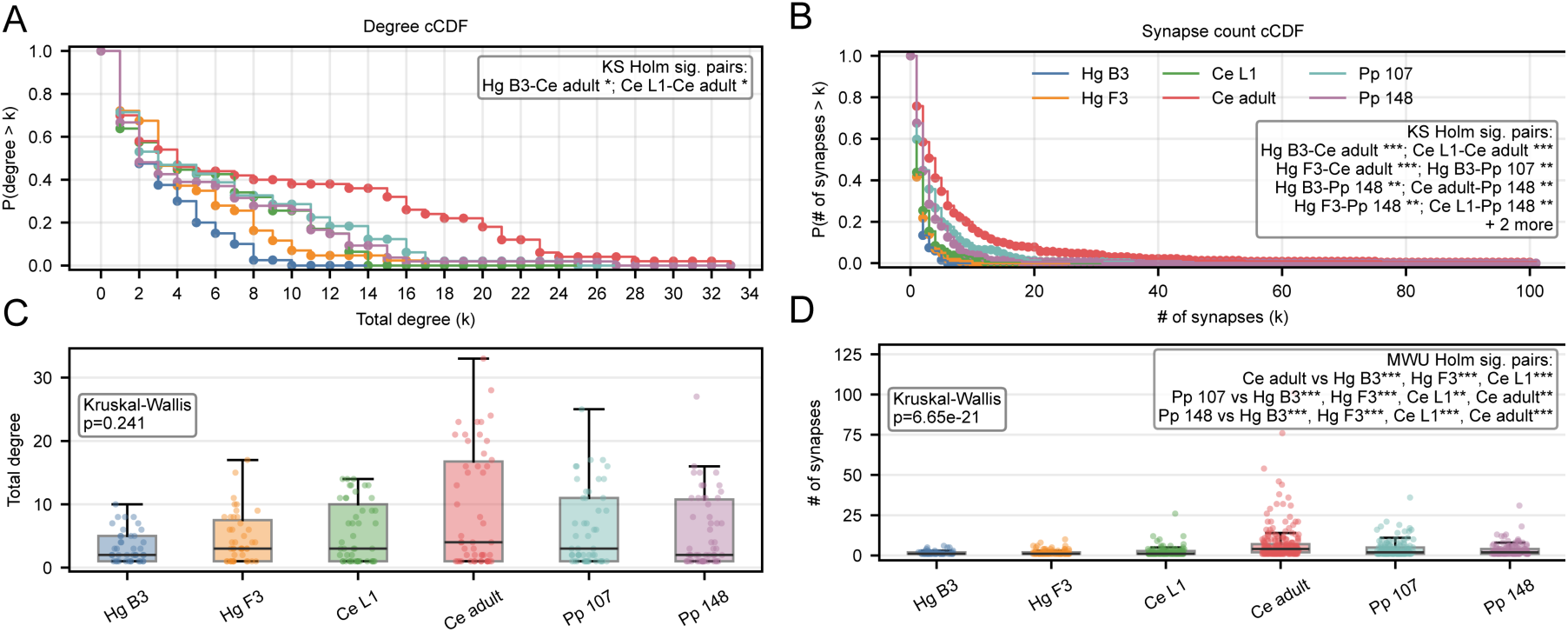
Degree and synaptic-weight distributions in esophageal connectomes. (A) Complementary cumulative distribution (cCDF) of total degree, P (degree > k). Distributions were compared pairwise using two-sample Kolmogorov–Smirnov (KS) tests with Holm correction. (B) cCDF of the number of synapses per connection, P(synapses > k). Distributions were compared pairwise using two-sample Kolmogorov–Smirnov (KS) tests with Holm correction. (C) Total degree per node by dataset. Differences across datasets were tested by Kruskal–Wallis. Brackets denote Holm-corrected pairwise comparisons. (D) Number of synapses per connection by dataset. Differences were tested using Kruskal–Wallis; brackets denote Holm-corrected pairwise Mann–Whitney U comparisons. Significance: *, p < 0.05; **, p < 0.01; ***, p < 0.001. Box, interquartile range; center line, median. Hg B3 and Hg F3, *H. glycines* J2; Ce L1, *C. elegans* late L1; Ce adult, *C. elegans* adult hermaphrodite; Pp 107 and Pp 148, *P. pacificus* adult hermaphrodite.

**Fig. S9.**
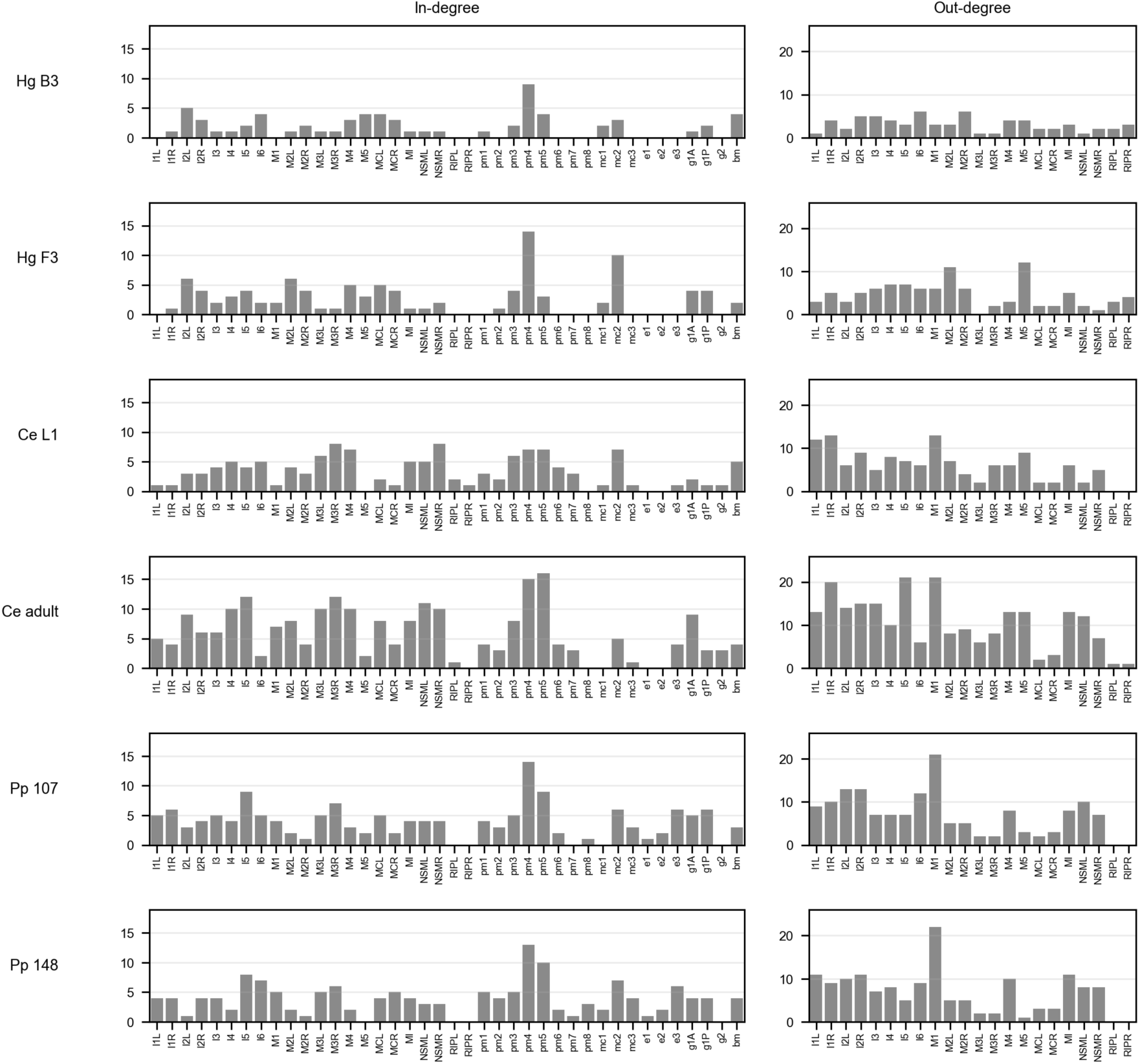
In-degree (left) and out-degree (right) profiles across esophageal connectomes. Hg B3 and Hg F3, *H. glycines* J2; Ce L1, *C. elegans* late L1; Ce adult, *C. elegans* adult hermaphrodite; Pp 107 and Pp 148, *P. pacificus* adult hermaphrodite. Non-neuronal cell name abbreviations follow **Table S1**.

**Table S1.**
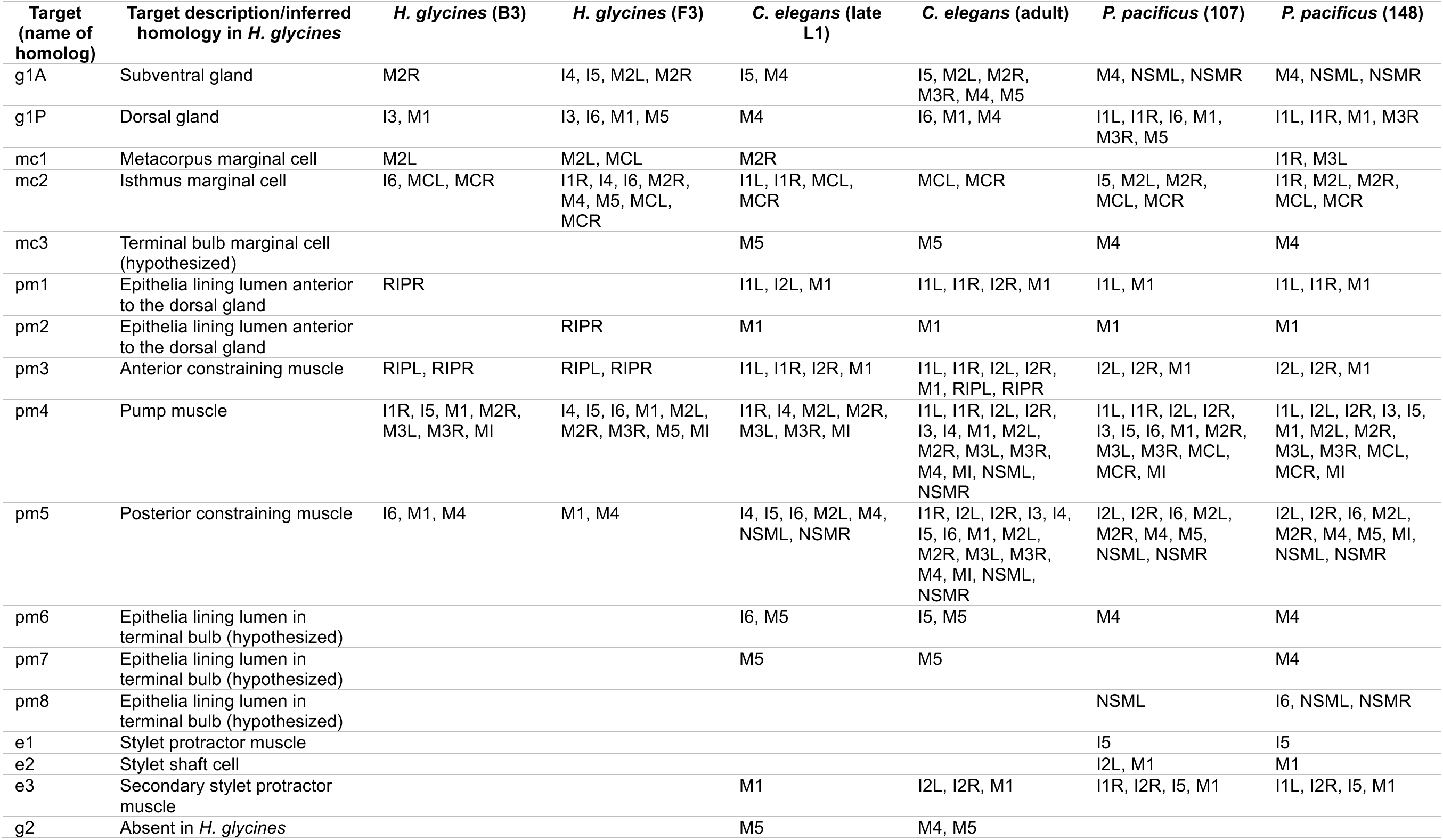
Neuronal inputs to esophageal target cell classes.

**Table S2.** Network properties of esophageal connectome datasets.

|  | *H. glycines* |  | *C. elegans* |  | *P. pacificus* (6) |  |
| --- | --- | --- | --- | --- | --- | --- |
|  | B3 | F3 | L1 | Adult (7) | Specimen 107 | Specimen 148 |
| Neuron, n<sup>*</sup> | 22 | 22 | 22 | 22 | 22 | 22 |
| Non-neuron, n | 18 | 21 | 25 | 28 | 27 | 32 |
| Connections | 67 | 101 | 130 | 231 | 154 | 151 |
| Network density | 0.043 | 0.056 | 0.060 | 0.094 | 0.065 | 0.053 |
| Average total degree | 3.4 | 4.7 | 5.5 | 9.2 | 6.3 | 5.6 |
| Diameter (undirected) | 7 | 6 | 6 | 5 | 4 | 4 |
| Average path length (undirected) | 3.32 | 2.84 | 2.79 | 2.38 | 2.45 | 2.57 |
|  | ×1.04<sup>¶</sup> | ×1.02 | ×1.04 | ×1.07 | ×1.04 | ×1.04 |
| Reciprocity | 0.21 | 0.20 | 0.15 | 0.35 | 0.18 | 0.13 |
|  | ×3.24 | ×2.46 | ×1.56 | ×1.43 | ×1.50 | ×1.33 |
| Transitivity | 0.07 | 0.11 | 0.14 | 0.29 | 0.16 | 0.11 |
|  | ×1.23 | ×1.37 | ×1.30 | ×1.19 | ×1.29 | ×1.11 |
| Average clustering coefficient | 0.08 | 0.18 | 0.26 | 0.39 | 0.34 | 0.28 |
|  | ×0.82 | ×1.04 | ×1.29 | ×0.93 | ×1.14 | ×1.14 |
| Largest SCC size<sup>†</sup> | 8 | 13 | 16 | 18 | 15 | 13 |
| SCC clustering coefficient | 0.43 | 0.46 | 0.45 | 0.74 | 0.62 | 0.41 |
|  | ×0.74 | ×1.09 | ×0.92 | ×1.07 | ×1.21 | ×0.84 |
| SCC path length | 2.29 | 2.46 | 2.16 | 1.63 | 2.04 | 2.07 |
|  | ×1.17 | ×1.10 | ×1.10 | ×1.03 | ×1.06 | ×1.02 |
| SCC small-worldness, S | 0.63 | 0.99 | 0.84 | 1.04 | 1.14 | 0.82 |
| Largest WCC<sup>‡</sup> | 40 | 43 | 47 | 50 | 49 | 54 |
| Synapse object | 107 | 178 | 218 | 1013 | 406 | 361 |
| Postsynaptic multiplicity (M/P), n<sup>§</sup> | 100/7 | 160/18 | 138/80 | 432/581 | 212/194 | 222/139 |
| Postsynaptic multiplicity (M/P), % | 93.5/6.5 | 89.9/10.1 | 63.3/36.7 | 42.6/57.4 | 52.2/47.8 | 61.5/38.5 |
| Total synaptic weight | 114 | 197 | 300 | 1619 | 611 | 511 |
| Total synaptic weight /connection | 1.70 | 1.95 | 2.31 | 7.01 | 3.97 | 3.38 |

\*20 esophageal neurons and two somatic neurons RIPL/R
<sup>†</sup>SCC, strongly connected component
<sup>‡</sup>WCC, weakly connected component
<sup>§</sup>M/P, monadic/polyadic postsynaptic multiplicity
<sup>¶</sup>Value after × are the ratio of the observed value to the mean of 500 randomized networks. Average path length, reciprocity, transitivity, average clustering coefficient were compared to averages over 500 degree-preserving random graphs. SCC clustering coefficient and SCC path length were compared to averages over 500 Erdős–Rényi random graphs.

**Table S3.** Observed normalized edit distance compared with degree-preserving randomized networks. For each dataset pair, the normalized edit distance between the observed networks was compared with a null distribution generated by degree-preserving randomization of both networks (n = 500). Observed, observed normalized edit distance; Expected, mean of the edit distance across the randomized ensemble; Std, standard deviation of the randomized ensemble. Hg B3 and Hg F3, *H. glycines* J2; Ce L1, *C. elegans* late L1; Ce adult, *C. elegans* adult hermaphrodite, Pp 107 and Pp 148, *P. pacificus* adult hermaphrodite. P values were adjusted using step-down min-P procedure.

| Dataset_1 | Dataset_2 | Observed | Expected | Std | Observed-expected<br>(Δ) | Z score | P value | Adjusted<br>P value |
| --- | --- | --- | --- | --- | --- | --- | --- | --- |
| Hg B3 | Hg F3 | 0.416 | 0.776 | 0.036 | -0.361 | -9.879 | 0.002 | 0.002 |
| Pp 107 | Pp 148 | 0.193 | 0.665 | 0.029 | -0.472 | -16.370 | 0.002 | 0.002 |
| Ce L1 | Ce adult | 0.423 | 0.662 | 0.024 | -0.239 | -10.116 | 0.002 | 0.002 |
| Ce L1 | Pp 107 | 0.623 | 0.736 | 0.029 | -0.113 | -3.883 | 0.002 | 0.002 |
| Ce L1 | Pp 148 | 0.624 | 0.747 | 0.028 | -0.123 | -4.464 | 0.002 | 0.002 |
| Ce adult | Pp 107 | 0.517 | 0.630 | 0.024 | -0.113 | -4.627 | 0.002 | 0.002 |
| Ce adult | Pp 148 | 0.544 | 0.665 | 0.023 | -0.120 | -5.176 | 0.002 | 0.002 |
| Hg B3 | Ce L1 | 0.792 | 0.864 | 0.028 | -0.071 | -2.566 | 0.010 | 0.086 |
| Hg B3 | Ce adult | 0.778 | 0.833 | 0.020 | -0.055 | -2.765 | 0.002 | 0.002 |
| Hg B3 | Pp 107 | 0.744 | 0.833 | 0.027 | -0.089 | -3.353 | 0.002 | 0.002 |
| Hg B3 | Pp 148 | 0.768 | 0.843 | 0.025 | -0.075 | -2.957 | 0.008 | 0.060 |
| Hg F3 | Ce L1 | 0.697 | 0.832 | 0.030 | -0.135 | -4.498 | 0.002 | 0.002 |
| Hg F3 | Ce adult | 0.674 | 0.771 | 0.021 | -0.097 | -4.549 | 0.002 | 0.002 |
| Hg F3 | Pp 107 | 0.714 | 0.800 | 0.026 | -0.086 | -3.251 | 0.002 | 0.002 |
| Hg F3 | Pp 148 | 0.752 | 0.822 | 0.026 | -0.070 | -2.660 | 0.012 | 0.112 |

**Table S4.** Raw centrality metrics (betweenness, PageRank, in-closeness, out-closeness), in- and out-degrees across six datasets. Hg B3 and Hg F3, *H. glycines*; Ce L1, *C. elegans* late L1; Ce adult, *C. elegans* adult hermaphrodite, Pp 107 and Pp 148, *P. pacificus* adult hermaphrodite.

| Metric | Node | B3 | F3 | Ce L1 | Ce adult | Pp 107 | Pp 148 |
| --- | --- | --- | --- | --- | --- | --- | --- |
| Betweenness | I1L | 0.000 | 0.000 | 0.043 | 0.016 | 0.018 | 0.022 |
| Betweenness | I1R | 0.005 | 0.092 | 0.023 | 0.006 | 0.025 | 0.022 |
| Betweenness | I2L | 0.014 | 0.073 | 0.092 | 0.040 | 0.026 | 0.005 |
| Betweenness | I2R | 0.052 | 0.127 | 0.021 | 0.019 | 0.040 | 0.021 |
| Betweenness | I3 | 0.020 | 0.027 | 0.052 | 0.013 | 0.043 | 0.012 |
| Betweenness | I4 | 0.023 | 0.004 | 0.067 | 0.014 | 0.054 | 0.046 |
| Betweenness | I5 | 0.003 | 0.054 | 0.049 | 0.099 | 0.057 | 0.024 |
| Betweenness | I6 | 0.075 | 0.017 | 0.021 | 0.002 | 0.070 | 0.053 |
| Betweenness | M1 | 0.000 | 0.018 | 0.078 | 0.065 | 0.043 | 0.046 |
| Betweenness | M2L | 0.022 | 0.143 | 0.020 | 0.004 | 0.007 | 0.003 |
| Betweenness | M2R | 0.064 | 0.019 | 0.018 | 0.002 | 0.001 | 0.001 |
| Betweenness | M3L | 0.001 | 0.000 | 0.003 | 0.004 | 0.002 | 0.005 |
| Betweenness | M3R | 0.001 | 0.000 | 0.057 | 0.020 | 0.004 | 0.003 |
| Betweenness | M4 | 0.032 | 0.016 | 0.080 | 0.032 | 0.031 | 0.026 |
| Betweenness | M5 | 0.076 | 0.098 | 0.000 | 0.031 | 0.001 | 0.000 |
| Betweenness | MCL | 0.013 | 0.014 | 0.000 | 0.011 | 0.003 | 0.002 |
| Betweenness | MCR | 0.014 | 0.003 | 0.000 | 0.004 | 0.001 | 0.004 |
| Betweenness | MI | 0.051 | 0.000 | 0.036 | 0.025 | 0.019 | 0.033 |
| Betweenness | NSML | 0.002 | 0.004 | 0.002 | 0.020 | 0.047 | 0.033 |
| Betweenness | NSMR | 0.022 | 0.014 | 0.030 | 0.012 | 0.038 | 0.038 |
| Betweenness | RIPL | 0.000 | 0.000 | 0.000 | 0.000 |  |  |
| Betweenness | RIPR | 0.000 | 0.000 | 0.000 | 0.000 |  |  |
| In-closeness | I1L | 0.000 | 0.000 | 0.145 | 0.285 | 0.206 | 0.174 |
| In-closeness | I1R | 0.089 | 0.170 | 0.115 | 0.243 | 0.214 | 0.182 |
| In-closeness | I2L | 0.211 | 0.259 | 0.198 | 0.330 | 0.180 | 0.121 |
| In-closeness | I2R | 0.125 | 0.250 | 0.169 | 0.285 | 0.206 | 0.182 |
| In-closeness | I3 | 0.079 | 0.174 | 0.209 | 0.275 | 0.222 | 0.182 |
| In-closeness | I4 | 0.071 | 0.160 | 0.254 | 0.344 | 0.222 | 0.143 |
| In-closeness | I5 | 0.096 | 0.221 | 0.215 | 0.375 | 0.288 | 0.261 |
| In-closeness | I6 | 0.178 | 0.153 | 0.237 | 0.229 | 0.222 | 0.235 |
| In-closeness | M1 | 0.000 | 0.156 | 0.145 | 0.306 | 0.199 | 0.182 |
| In-closeness | M2L | 0.129 | 0.214 | 0.222 | 0.318 | 0.175 | 0.133 |
| In-closeness | M2R | 0.093 | 0.179 | 0.203 | 0.243 | 0.144 | 0.114 |
| In-closeness | M3L | 0.129 | 0.033 | 0.237 | 0.344 | 0.221 | 0.204 |
| In-closeness | M3R | 0.123 | 0.150 | 0.284 | 0.375 | 0.276 | 0.213 |
| In-closeness | M4 | 0.152 | 0.294 | 0.274 | 0.330 | 0.207 | 0.156 |
| In-closeness | M5 | 0.164 | 0.183 | 0.000 | 0.229 | 0.175 | 0.000 |
| In-closeness | MCL | 0.251 | 0.305 | 0.132 | 0.319 | 0.269 | 0.181 |
| In-closeness | MCR | 0.221 | 0.244 | 0.102 | 0.264 | 0.189 | 0.231 |
| In-closeness | MI | 0.107 | 0.136 | 0.222 | 0.306 | 0.214 | 0.182 |
| In-closeness | NSML | 0.108 | 0.153 | 0.243 | 0.344 | 0.222 | 0.174 |
| In-closeness | NSMR | 0.084 | 0.231 | 0.274 | 0.344 | 0.214 | 0.167 |
| In-closeness | RIPL | 0.000 | 0.000 | 0.146 | 0.206 |  |  |
| In-closeness | RIPR | 0.000 | 0.000 | 0.122 | 0.000 |  |  |
| In-degree | I1L | 0.000 | 0.000 | 1.000 | 5.000 | 5.000 | 4.000 |
| In-degree | I1R | 1.000 | 1.000 | 1.000 | 4.000 | 6.000 | 4.000 |
| In-degree | I2L | 5.000 | 6.000 | 3.000 | 9.000 | 3.000 | 1.000 |
| In-degree | I2R | 3.000 | 4.000 | 3.000 | 6.000 | 4.000 | 4.000 |
| In-degree | I3 | 1.000 | 2.000 | 4.000 | 6.000 | 5.000 | 4.000 |
| In-degree | I4 | 1.000 | 2.000 | 5.000 | 10.000 | 4.000 | 2.000 |
| In-degree | I5 | 2.000 | 4.000 | 3.000 | 12.000 | 9.000 | 8.000 |
| In-degree | I6 | 4.000 | 2.000 | 5.000 | 2.000 | 5.000 | 7.000 |
| In-degree | M1 | 0.000 | 2.000 | 1.000 | 7.000 | 4.000 | 5.000 |
| In-degree | M2L | 1.000 | 5.000 | 4.000 | 8.000 | 2.000 | 2.000 |
| In-degree | M2R | 2.000 | 4.000 | 3.000 | 4.000 | 1.000 | 1.000 |
| In-degree | M3L | 1.000 | 1.000 | 6.000 | 10.000 | 5.000 | 5.000 |
| In-degree | M3R | 1.000 | 1.000 | 8.000 | 12.000 | 7.000 | 6.000 |
| In-degree | M4 | 3.000 | 5.000 | 7.000 | 9.000 | 3.000 | 2.000 |
| In-degree | M5 | 4.000 | 3.000 | 0.000 | 2.000 | 2.000 | 0.000 |
| In-degree | MCL | 4.000 | 5.000 | 2.000 | 8.000 | 5.000 | 4.000 |
| In-degree | MCR | 3.000 | 4.000 | 1.000 | 4.000 | 2.000 | 5.000 |
| In-degree | MI | 1.000 | 1.000 | 5.000 | 8.000 | 4.000 | 4.000 |
| In-degree | NSML | 1.000 | 1.000 | 5.000 | 10.000 | 4.000 | 3.000 |
| In-degree | NSMR | 1.000 | 2.000 | 8.000 | 10.000 | 4.000 | 3.000 |
| In-degree | RIPL | 0.000 | 0.000 | 2.000 | 1.000 |  |  |
| In-degree | RIPR | 0.000 | 0.000 | 1.000 | 0.000 |  |  |
| In-degree | bm | 4.000 | 2.000 | 5.000 | 4.000 | 3.000 | 4.000 |
| In-degree | e1 |  |  |  |  | 1.000 | 1.000 |
| In-degree | e2 |  |  |  |  | 2.000 | 1.000 |
| In-degree | e3 |  |  | 1.000 | 3.000 | 4.000 | 4.000 |
| In-degree | g1A | 1.000 | 4.000 | 2.000 | 6.000 | 3.000 | 3.000 |
| In-degree | g1P | 2.000 | 4.000 | 1.000 | 3.000 | 6.000 | 4.000 |
| In-degree | g2 |  |  | 1.000 | 2.000 |  |  |
| In-degree | mc1 | 1.000 | 2.000 | 1.000 |  |  | 2.000 |
| In-degree | mc2 | 3.000 | 8.000 | 4.000 | 2.000 | 5.000 | 5.000 |
| In-degree | mc3 |  |  | 1.000 | 1.000 | 1.000 | 2.000 |
| In-degree | pm1 | 1.000 |  | 3.000 | 4.000 | 2.000 | 3.000 |
| In-degree | pm2 |  | 1.000 | 1.000 | 1.000 | 1.000 | 1.000 |
| In-degree | pm3 | 2.000 | 2.000 | 4.000 | 7.000 | 3.000 | 3.000 |
| In-degree | pm4 | 7.000 | 9.000 | 7.000 | 15.000 | 14.000 | 13.000 |
| In-degree | pm5 | 3.000 | 2.000 | 7.000 | 16.000 | 9.000 | 10.000 |
| In-degree | pm6 |  |  | 2.000 | 2.000 | 1.000 | 1.000 |
| In-degree | pm7 |  |  | 1.000 | 1.000 |  | 1.000 |
| In-degree | pm8 |  |  |  |  | 1.000 | 3.000 |
| Out-closeness | I1L | 0.076 | 0.223 | 0.438 | 0.541 | 0.447 | 0.500 |
| Out-closeness | I1R | 0.263 | 0.384 | 0.451 | 0.648 | 0.507 | 0.431 |
| Out-closeness | I2L | 0.089 | 0.246 | 0.418 | 0.579 | 0.586 | 0.508 |
| Out-closeness | I2R | 0.270 | 0.315 | 0.356 | 0.601 | 0.586 | 0.508 |
| Out-closeness | I3 | 0.194 | 0.315 | 0.374 | 0.601 | 0.493 | 0.473 |
| Out-closeness | I4 | 0.349 | 0.376 | 0.384 | 0.508 | 0.479 | 0.500 |
| Out-closeness | I5 | 0.033 | 0.305 | 0.401 | 0.688 | 0.405 | 0.143 |
| Out-closeness | I6 | 0.295 | 0.337 | 0.284 | 0.429 | 0.540 | 0.442 |
| Out-closeness | M1 | 0.100 | 0.320 | 0.499 | 0.648 | 0.548 | 0.486 |
| Out-closeness | M2L | 0.075 | 0.480 | 0.360 | 0.465 | 0.391 | 0.354 |
| Out-closeness | M2R | 0.325 | 0.349 | 0.293 | 0.479 | 0.358 | 0.354 |
| Out-closeness | M3L | 0.033 | 0.000 | 0.245 | 0.423 | 0.066 | 0.056 |
| Out-closeness | M3R | 0.033 | 0.033 | 0.313 | 0.459 | 0.059 | 0.056 |
| Out-closeness | M4 | 0.267 | 0.067 | 0.331 | 0.524 | 0.118 | 0.178 |
| Out-closeness | M5 | 0.310 | 0.492 | 0.300 | 0.508 | 0.362 | 0.037 |
| Out-closeness | MCL | 0.076 | 0.067 | 0.028 | 0.029 | 0.059 | 0.056 |
| Out-closeness | MCR | 0.067 | 0.033 | 0.028 | 0.029 | 0.059 | 0.056 |
| Out-closeness | MI | 0.305 | 0.376 | 0.331 | 0.560 | 0.459 | 0.549 |
| Out-closeness | NSML | 0.033 | 0.315 | 0.056 | 0.533 | 0.500 | 0.473 |
| Out-closeness | NSMR | 0.150 | 0.033 | 0.306 | 0.435 | 0.447 | 0.473 |
| Out-closeness | RIPL | 0.100 | 0.217 | 0.000 | 0.029 |  |  |
| Out-closeness | RIPR | 0.248 | 0.276 | 0.000 | 0.029 |  |  |
| Out-degree | I1L | 1.000 | 3.000 | 11.000 | 13.000 | 9.000 | 11.000 |
| Out-degree | I1R | 4.000 | 5.000 | 12.000 | 19.000 | 10.000 | 9.000 |
| Out-degree | I2L | 2.000 | 3.000 | 6.000 | 14.000 | 13.000 | 10.000 |
| Out-degree | I2R | 5.000 | 5.000 | 9.000 | 15.000 | 13.000 | 11.000 |
| Out-degree | I3 | 5.000 | 6.000 | 5.000 | 15.000 | 7.000 | 7.000 |
| Out-degree | I4 | 4.000 | 6.000 | 8.000 | 10.000 | 7.000 | 8.000 |
| Out-degree | I5 | 1.000 | 5.000 | 6.000 | 20.000 | 7.000 | 5.000 |
| Out-degree | I6 | 6.000 | 5.000 | 6.000 | 6.000 | 12.000 | 9.000 |
| Out-degree | M1 | 3.000 | 6.000 | 11.000 | 18.000 | 13.000 | 12.000 |
| Out-degree | M2L | 2.000 | 10.000 | 7.000 | 8.000 | 5.000 | 5.000 |
| Out-degree | M2R | 6.000 | 6.000 | 4.000 | 9.000 | 5.000 | 5.000 |
| Out-degree | M3L | 1.000 | 0.000 | 2.000 | 6.000 | 2.000 | 2.000 |
| Out-degree | M3R | 1.000 | 1.000 | 6.000 | 8.000 | 2.000 | 2.000 |
| Out-degree | M4 | 3.000 | 2.000 | 6.000 | 11.000 | 4.000 | 6.000 |
| Out-degree | M5 | 4.000 | 11.000 | 5.000 | 7.000 | 3.000 | 1.000 |
| Out-degree | MCL | 2.000 | 2.000 | 1.000 | 1.000 | 2.000 | 2.000 |
| Out-degree | MCR | 2.000 | 1.000 | 1.000 | 1.000 | 2.000 | 2.000 |
| Out-degree | MI | 3.000 | 4.000 | 6.000 | 13.000 | 8.000 | 11.000 |
| Out-degree | NSML | 1.000 | 2.000 | 2.000 | 11.000 | 9.000 | 8.000 |
| Out-degree | NSMR | 2.000 | 1.000 | 5.000 | 7.000 | 7.000 | 8.000 |
| Out-degree | RIPL | 2.000 | 2.000 | 0.000 | 1.000 |  |  |
| Out-degree | RIPR | 3.000 | 3.000 | 0.000 | 1.000 |  |  |
| PageRank | I1L | 0.015 | 0.016 | 0.019 | 0.025 | 0.025 | 0.023 |
| PageRank | I1R | 0.021 | 0.023 | 0.016 | 0.019 | 0.028 | 0.023 |
| PageRank | I2L | 0.067 | 0.042 | 0.023 | 0.032 | 0.021 | 0.017 |
| PageRank | I2R | 0.034 | 0.042 | 0.020 | 0.025 | 0.026 | 0.024 |
| PageRank | I3 | 0.023 | 0.025 | 0.029 | 0.024 | 0.030 | 0.023 |
| PageRank | I4 | 0.019 | 0.024 | 0.034 | 0.036 | 0.028 | 0.020 |

| **Metric** | **Node** | **B3** | **F3** | **Ce L1** | **Ce adult** | **Pp 107** | **Pp 148** |
| --- | --- | --- | --- | --- | --- | --- | --- |
| PageRank | I5 | 0.023 | 0.030 | 0.025 | 0.037 | 0.040 | 0.034 |
| PageRank | I6 | 0.038 | 0.021 | 0.035 | 0.020 | 0.030 | 0.030 |
| PageRank | M1 | 0.015 | 0.022 | 0.018 | 0.025 | 0.025 | 0.025 |
| PageRank | M2L | 0.020 | 0.038 | 0.027 | 0.029 | 0.020 | 0.019 |
| PageRank | M2R | 0.026 | 0.029 | 0.027 | 0.019 | 0.019 | 0.018 |
| PageRank | M3L | 0.020 | 0.021 | 0.031 | 0.034 | 0.027 | 0.026 |
| PageRank | M3R | 0.024 | 0.018 | 0.046 | 0.043 | 0.034 | 0.027 |
| PageRank | M4 | 0.035 | 0.036 | 0.043 | 0.032 | 0.022 | 0.019 |
| PageRank | M5 | 0.041 | 0.027 | 0.014 | 0.016 | 0.020 | 0.015 |
| PageRank | MCL | 0.057 | 0.047 | 0.017 | 0.033 | 0.040 | 0.023 |
| PageRank | MCR | 0.053 | 0.043 | 0.015 | 0.019 | 0.023 | 0.030 |
| PageRank | MI | 0.024 | 0.018 | 0.038 | 0.033 | 0.027 | 0.024 |
| PageRank | NSML | 0.024 | 0.020 | 0.030 | 0.035 | 0.026 | 0.023 |
| PageRank | NSMR | 0.019 | 0.028 | 0.046 | 0.033 | 0.025 | 0.022 |
| PageRank | RIPL | 0.015 | 0.016 | 0.018 | 0.015 |  |  |
| PageRank | RIPR | 0.015 | 0.016 | 0.016 | 0.014 |  |  |
| PageRank | bm | 0.069 | 0.048 | 0.051 | 0.027 | 0.023 | 0.029 |
| PageRank | e1 |  |  |  |  | 0.020 | 0.021 |
| PageRank | e2 |  |  |  |  | 0.019 | 0.017 |
| PageRank | e3 |  |  | 0.016 | 0.018 | 0.026 | 0.026 |
| PageRank | g1A | 0.019 | 0.032 | 0.024 | 0.029 | 0.026 | 0.023 |
| PageRank | g1P | 0.023 | 0.029 | 0.020 | 0.020 | 0.044 | 0.033 |
| PageRank | g2 |  |  | 0.017 | 0.018 |  |  |
| PageRank | mc1 | 0.024 | 0.039 | 0.020 |  |  | 0.028 |
| PageRank | mc2 | 0.067 | 0.105 | 0.044 | 0.058 | 0.054 | 0.046 |
| PageRank | mc3 |  |  | 0.017 | 0.016 | 0.020 | 0.060 |
| PageRank | pm1 | 0.019 |  | 0.020 | 0.019 | 0.020 | 0.021 |
| PageRank | pm2 |  | 0.021 | 0.016 | 0.015 | 0.017 | 0.017 |
| PageRank | pm3 | 0.026 | 0.028 | 0.020 | 0.045 | 0.020 | 0.020 |
| PageRank | pm4 | 0.091 | 0.061 | 0.053 | 0.051 | 0.094 | 0.084 |
| PageRank | pm5 | 0.035 | 0.034 | 0.056 | 0.054 | 0.043 | 0.050 |
| PageRank | pm6 |  |  | 0.022 | 0.017 | 0.020 | 0.018 |
| PageRank | pm7 |  |  | 0.017 | 0.016 |  | 0.018 |
| PageRank | pm8 |  |  |  |  | 0.018 | 0.023 |

**Table S5.**
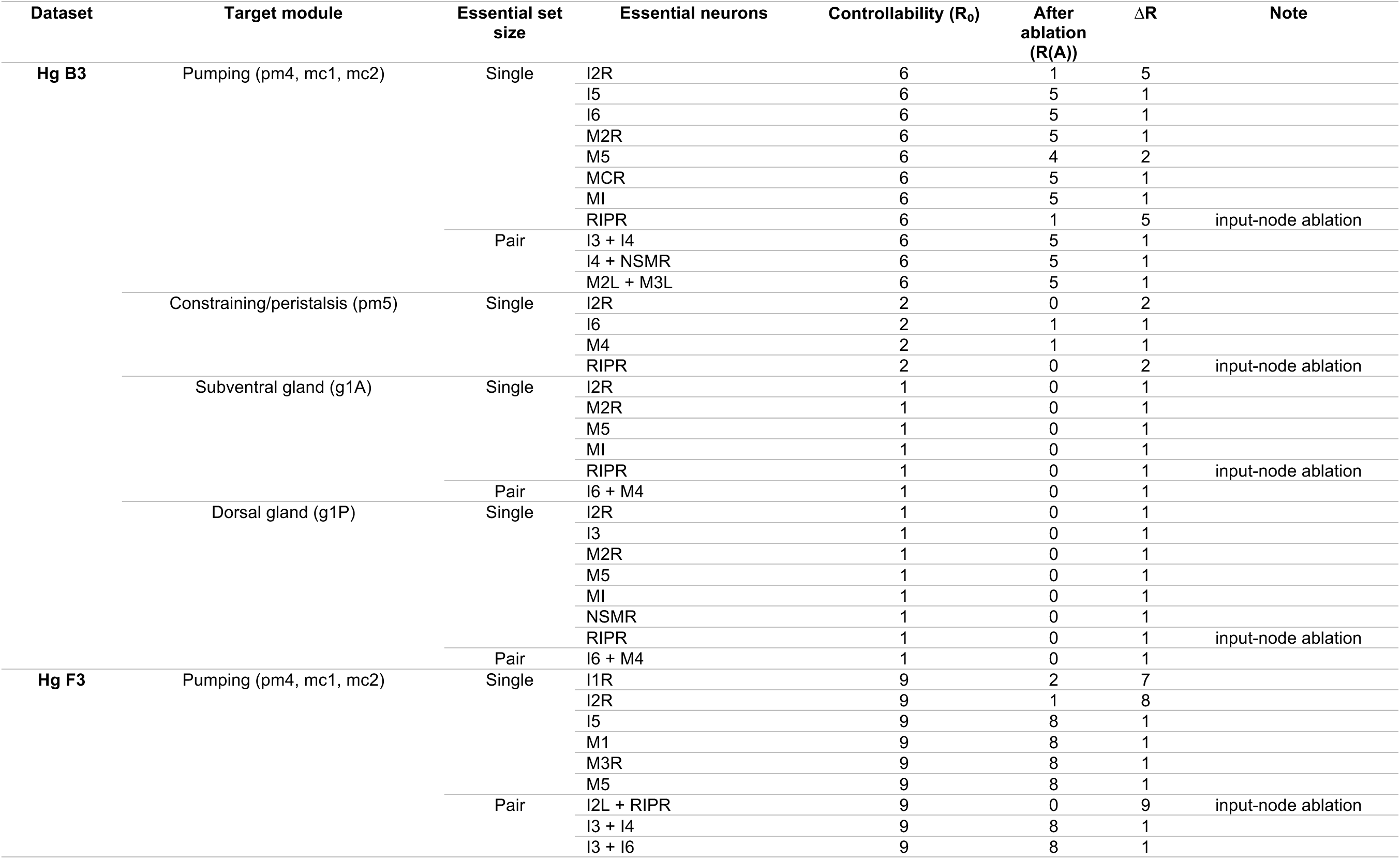
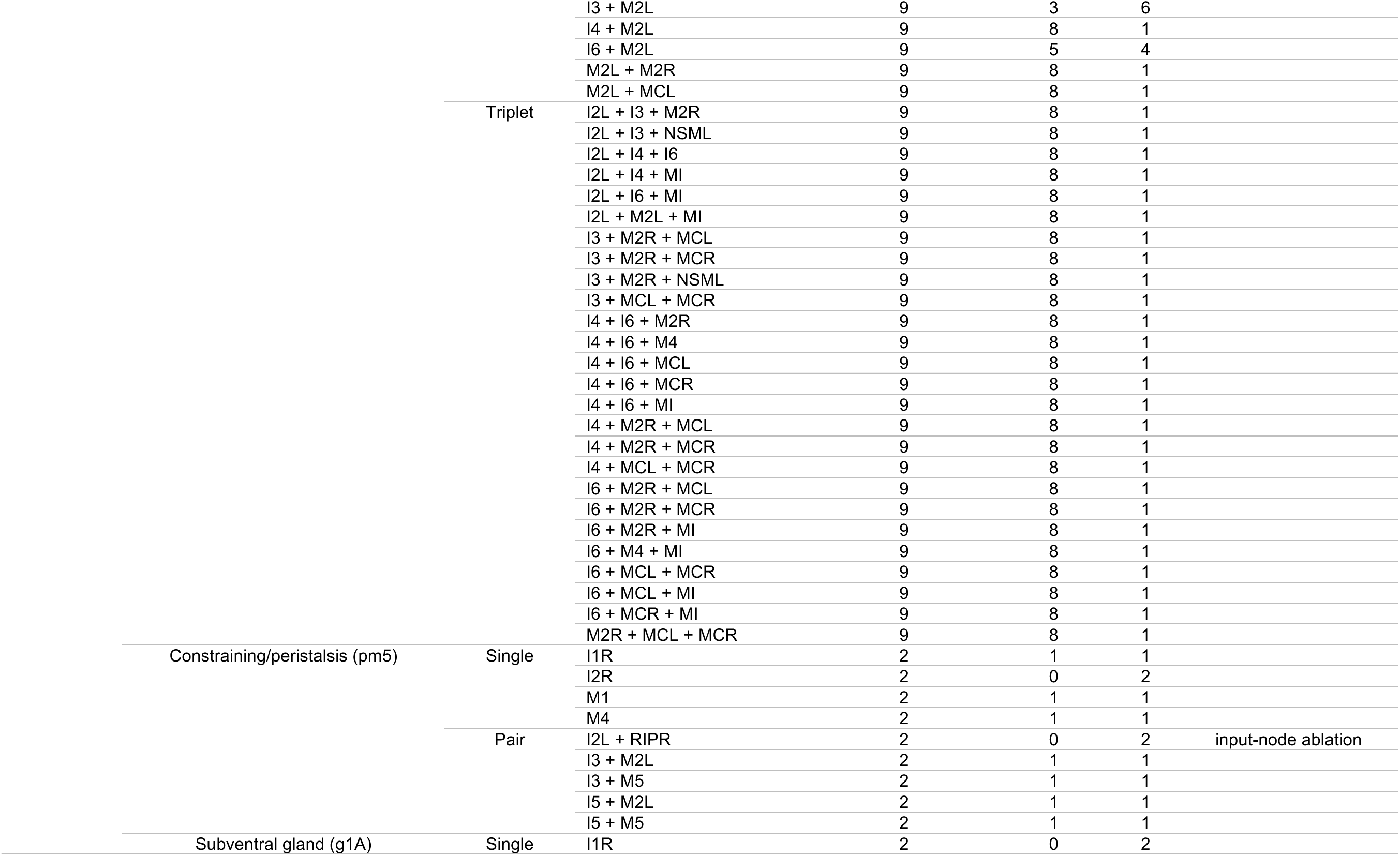
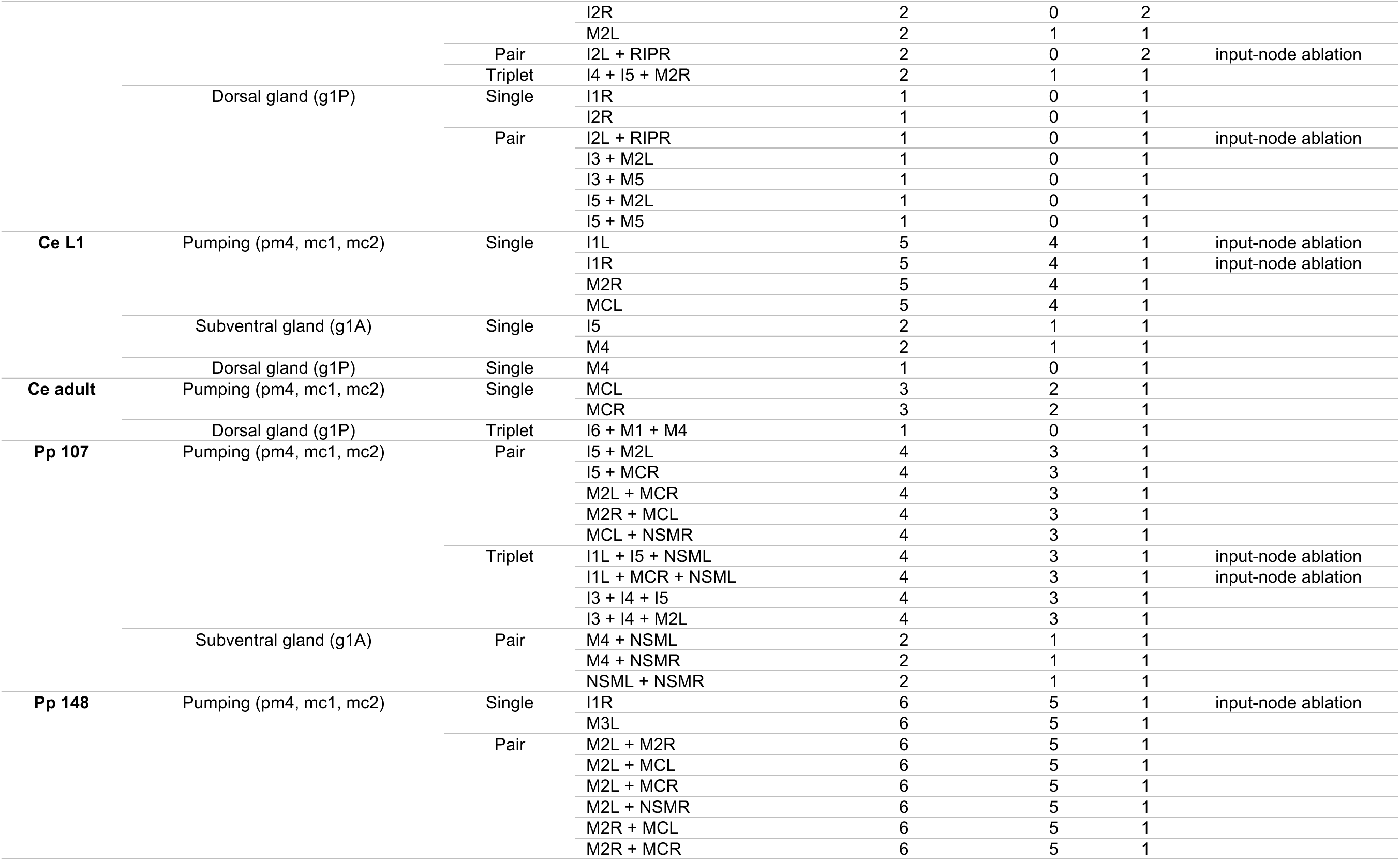
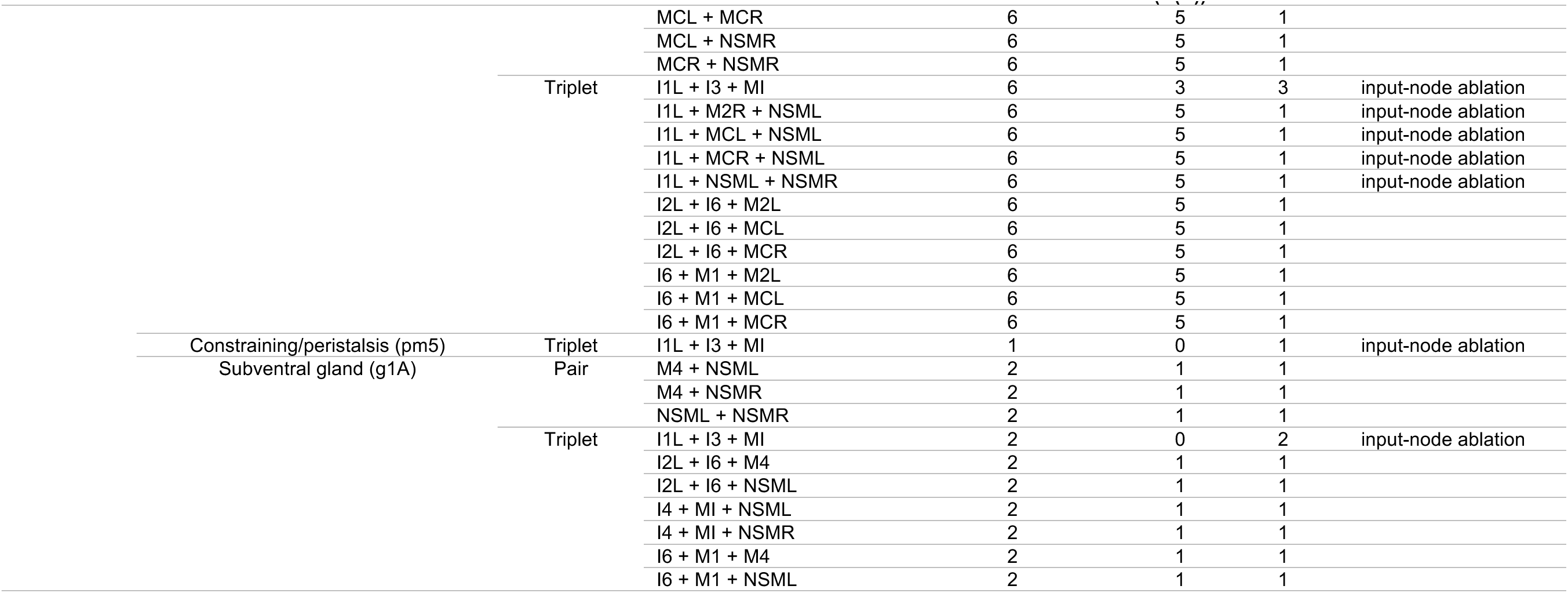
Synthetic ablation analysis of the esophageal connectome. Single essential neurons and synthetic essential pairs and triplets for each target module in *H. glycines* and free-living datasets. R0, baseline controllability score; R(A), recalculated score after ablation; ΔR, R_0_ – R(A).

**Dataset S1 (separate file).** Three-dimensional renderings of the esophageal neurons and the anterior processes of RIPL/R in *H. glycines* dataset F3. Each page shows a single neuron (red) in five orientations, from top to bottom: left lateral, dorsal, ventral, and two oblique lateral views taken from anterodorsal and posterodorsal positions. As landmarks, the esophageal lumen and the outer surface of the esophagus are shown in green and light green, respectively.

